# Effects of haemosporidian co-infection and parasitemia on reproductive strategies in a polymorphic species

**DOI:** 10.1101/2022.03.28.486032

**Authors:** Zoé Delefortrie, Hugo F. Gante, Oscar Gordo, Kristopher R. Schwab, Rusty A. Gonser

**Affiliations:** Department of Biology, Indiana State University, 47809 Terre Haute, IN, USA; The Center for Genomic Advocacy, Indiana State University, 47809 Terre Haute, IN, USA; Department of Biology, KU Leuven, 3000 Leuven, Belgium; Royal Museum for Central Africa, 3080 Tervuren, Belgium; Catalan Ornithological Institute, 08019 Barcelona, Spain

**Keywords:** Haemosporidians, co-infection, parasitemia, polymorphic species, terminal investment, reproductive strategies

## Abstract

Host-parasite interactions play a significant role in evolution. Parasite infections impose fitness costs that can trigger trade-off responses in host reproductive strategies. Individuals can invest limited resources in parasite defense, such as stimulating the immune system, or perform terminal reproductive investment. Here, we investigated how haemosporidian co-infection and parasitemia influence different reproductive strategies in a polymorphic bird species (the white-throated sparrow, *Zonotrichia albicollis*). We provided an account of the impacts of co-infection and parasitemia on host reproductive success and reproductive strategies in response to parasite infection. We tested the effect of co-infection and parasitemia on reproductive success (fledgling number, extra-pair paternity, ratio social/genetic offspring) and the effect of parental infection on nestling quality (mean nestling growth rate and body condition). We found that co-infection increases reproductive extra-pair paternity and nestling mean body condition. Parental high parasitemia positively impacts the ratio of genetic offspring belonging to the social father and has mixed results on nestling quality. We observed that co-infection in females and parasitemia in males might trigger a switch in reproductive strategy towards a terminal investment for co-infected individuals. In contrast, high parasitemia in females negatively impacted their offspring fitness, possibly due to the reallocation of resources for fighting the infection.

## INTRODUCTION

Host-parasite interactions play a significant role in the evolution and ecology of organisms (Cuevas et al. 2021, Poulin 2011). Parasites inflict substantial fitness costs by impacting host body condition, survival, or reproductive success (Van Rooyen et al. 2013). Hosts may alter their reproductive strategies in response to the handicap imposed by parasites (Agnew et al. 2000, Kulma et al. 2014, Schwanz 2008). One strategy can be a reallocation of resources for reproduction to the immune system for fighting against the parasite to increase resistance and chances of survival at the cost of reduced reproductive success (Nordling et al. 1998, Sheldon and Verhulst 1996). An alternative strategy could be an overinvestment in current reproduction to maximize immediate fitness, possibly at the expense of own survival (Agnew et al. 2000). Thus, host species may show various strategies depending on their reproductive mode and infection severity. In this evolutionary scenario, behaviorally polymorphic species are excellent study models to understand parasite-mediated life-history trade-offs.

The avian malaria model has been extensively used to study host-parasite co-evolution, including infection costs (Fallon et al. 2003, Mukhin et al. 2016, Schoenle et al. 2017, Zehtindjiev et al. 2008) and life history trade-offs (Gilman et al. 2007, Ishtiaq et al. 2017, Karell et al. 2007, Knowles et al. 2011, MacDougall-Shackleton et al. 2002). Three genera of the order Haemosporida – *Plasmodium, Haemoproteus*, and *Leucocytozoon* – cause avian malaria and are present in almost every bird community in the world (LaPointe et al. 2012, Schoenle et al. 2017, Valkiūnas 2004). For instance, depending on species and region, their prevalence varies from 50% to 80% in North American birds (Reinoso-Pérez et al. 2016).

Usually, multiple strains or even different species of these malaria parasites infect hosts in wild populations (Palinauskas et al. 2016, Pigeault et al. 2020, Reinoso-Pérez et al. 2020, Valkiūnas et al. 2003). Furthermore, co-infecting parasites interact not only with the host but with each other (Bose et al. 2016). Thus, host-parasite and parasite-parasite interactions lead to variation in parasitemia, impacting virulence and infection dynamics (Bose et al. 2016, Van Rooyen et al. 2013). While the effect of malaria prevalence on host fitness has been widely investigated (Cuevas et al. 2021, De La Torre et al. 2020, Figuerola et al. 1999, Jacobs et al. 2015), less often those studies integrated the costs of co-infection on host reproductive success (Marzal et al. 2008, Van Rooyen et al. 2013, Norte et al. 2009, Pigeault et al. 2020, Pigeault et al. 2018).

Infection costs depend not only on prevalence (presence of the parasite) but also on parasitemia (amount of parasites within a host) (Schoenle et al. 2017). Knowles et al. (2011) suggested that parasitemia might be more informative about host-parasite interactions following an infection, such as host immune response to parasite invasion. In contrast, prevalence provides information about the degree of exposure to parasite vectors. Interestingly, when looking at the effect of parasite co-infection and parasitemia on host reproductive success, we often find mixed results across life-history traits, such as clutch size (Marzal et al. 2008), and hatching (Marzal et al. 2005, Knowles et al. 2010, Merino et al. 2000) and fledgling success (Marzal et al. 2008, Norte et al. 2009, Pigeault et al. 2020). Co-infections have been observed to positively affect reproductive success by increasing clutch size (Marzal et al. 2008) and increasing the number of fledglings (Marzal et al. 2008, Norte et al. 2009, Pigeault et al. 2020). On the other hand, high parasitemia is detrimental to host fitness by reducing clutch size, hatching success, and fledgling success (Marzal et al. 2005, Knowles et al. 2010, Merino et al. 2000). In the case of extra-pair paternity, Jacobs et al. (2015) found that males infected by avian malaria are not likely to lose paternity in their own nest. Still, they are less likely to gain extra-pair paternity in another nest. But males with a higher level infection are more likely to acquire extra-pair paternity and have higher chances to get co-infected with other parasites (Badás et al. 2020). Extra-pair paternity improves male reproductive success and benefits females (Forstmeier et al. 2014, Vedder et al. 2011). Under high parasite affection, females would seek extra-pair copulation to enhance offspring genetic diversity, which may strengthen offspring immune system and improve female reproductive success (Forstmeier et al. 2014, Podmokła et al. 2015).

Parental infections indirectly affect offspring quality by affecting parental performance during parental care stages. The impact on parental performance would be especially relevant for females, which usually have a more prominent role in nestling care (Merino et al. 2000, Pigeault et al. 2020). Depending on parasitemia level, females may apply different parental strategies. For example, heavily infected females would invest more in parental care to compensate for the reduced chances of future reproduction, performing a terminal investment (Schwanz 2008, Part et al. 1992, Kulma et al. 2014). In comparison, individuals with a medium or low infection would invest more resources in their immune system and thus reducing their reproductive output to survive the disease (Reaney and Knell 2010). Therefore, infected individuals have a trade-off between actual and future reproductive success based on their survival prospects until the next reproductive cycle (Agnew et al. 2000).

Polymorphic species are species in which individuals present different morphs determined genetically (Roulin 2004). Each morph may have different parasite prevalence and parasitemia (Boyd et al. 2018, Galeotti and Sacchi 2003, Gangoso et al. 2016, Karell et al. 2011, Lei et al. 2014), as they have different strategies for resource allocation to reproduction and immunity (Boyd et al. 2018, Galeotti and Sacchi 2003, Karell et al. 2017). For example, in tawny owls (*Strix aluco*), only rufous individuals with no infection or low parasitemia are able to reproduce, while the parasite burden does not affect the reproductive success of grey birds (Galeotti and Sacchi 2003). The low impact of parasite infection on the grey morph could result from better exploitation of parasitic reduction with high immune response linked to a reduction in telomere length (Karell et al. 2017, Karell et al. 2011). In white-throated sparrows (*Zonotrichia albicollis*), tan individuals show higher parasitemia than white ones but no differences in prevalence in artificially infected birds with *Plasmodium* (Boyd et al. 2018). The differences in parasitemia were suggested to be due to differences in life history and potentially linked to differences in parasite resistance (Boyd et al. 2018). No one has yet tested whether the white-throated sparrow alternative strategies are sufficient to fight the parasite infection costs.

We studied host-parasite interactions between malaria (*Plasmodium* and *Leucocytozoon*) and white-throated sparrows. The white-throated sparrow is a North American passerine with a chromosomal polymorphism (Thorneycroft 1966). This chromosomal polymorphism is due to a rearrangement on chromosome 2 linked to a phenotypic and behavioral polymorphism with differences in plumage color (white and tan individuals), territorial defense, and reproductive strategy (Lowther 1962, Thorneycroft 1966, Thorneycroft 1975, Tuttle 2003). White morph tends to be more aggressive and provide less parental care than tan morph (Tuttle 2003). In addition, white males attempt more extra-pair copulations leading to higher extra-pair paternity than tan males (Tuttle 2003). However, the proportion of both morphs is relatively stable due to disassortative mating, where tan males mate with white females (T×W) and white males pair with tan females (W×T) (Tuttle et al. 2016). The behavioral differences between morphs imply antagonistic reproductive strategies for each pair type, giving similar fitness to each morph and consequently keeping the equilibrium between the morphs in a population (Grunst et al. 2019b).

We investigated the impact of co-infection and parasitemia on adult body condition, reproductive success (number of fledglings, chance of extra-pair paternity in a nest, and the ratio of genetic offspring belonging to the social father), and how parental infection impacts nestling condition. We predicted: 1) body condition will be worse in those individuals infected by two haemosporidian species and having a higher parasitemia (De La Torre et al. 2020); 2) individuals with higher parasitemia will produce more fledglings in a single season if they performed a terminal investment strategy; 3) females with infected partners will show higher extra-pair paternity, as they would actively seek healthier males. Furthermore, when investigating the effect of parental infection on nestlings, we predicted that co-infected parents with high parasitemia will have nestlings in better body condition and with faster growth rates.

## MATERIALS AND METHODS

### Study site and data collection

We studied a population of white-throated sparrows between 2002 and 2019 near the Cranberry Lake Biological Station in the Adirondack Mountains (44.15ºN - 74.78ºW, New York, USA). The study site consists of bog, forest, and pond, prime habitats for white-throated sparrow reproduction. From 2002 to 2019, 952 pairs have been studied with a mean population density of 0.39 ± 0.1 pairs per Ha over a mean of 139.32 ± 39.5 Ha surveyed every year. Following Tuttle (2003), behavioral data were collected on territories with stable and delimited borders (see Supplementary information for details). Adults were identified by their unique combinations of color rings. Based on adult behavior, pair’s nests were found, and their reproduction status was established (nest building, laying, incubation, nestling rearing). For every pair’s nest, morphometric measurements were recorded such as tarsus length to the nearest 0.01mm with digital calipers and body mass to the nearest 0.5 g with Pesola^®^ spring scale from hatching day till day 6. On day 6 post-hatching, nestlings and adults were banded with an individual color combination and a U.S. Fish & Wildlife Service aluminum band. A small blood sample was collected from the brachial vein (∼ 80 μl) for each nestling. Fledging success was determined by the number of chicks banded at the nest at that moment. Additionally, adult blood samples were collected during their 1^st^ clutch when they had nestling with ages between 4 to 10 days. Blood samples were preserved in Longmire buffer (Longmire et al. 1992) until we extracted the DNA with DNeasy^®^ Blood & tissue extraction kit (Qiagen, Germantown, MD, USA).

Individual body condition was calculated based on a scaled mass index (Peig and Green 2009). For the offspring, the body condition was averaged per nest. In addition, offspring growth rate was calculated based on the slope between weight and day and took the average of the nest (Grunst et al. 2019a).

### Adult morph and parentage analysis

Adult birds were sexed based on their phenotype and behavior: males had cloacal protuberance and displayed territory defense, while females showed brood patch. Adult morph (white or tan) was confirmed by genetic analyses (Michopoulos et al. 2007). To assess reproductive success, the number of genetic offspring (paternity) was determined and adjusted with the allocation of extra-pair paternity (EPP) by following the protocol of Grunst et al. (2019b). In our statistical analysis, EPP was qualified present in a nest when the social fathers were not the genetic father of one or more of the nestlings. The EPP variable was used as a binomial variable (EPP absent: 0 / EPP present: 1). In addition, a new metric was implemented – the ratio of genetic offspring belonging to the social father (G_o_/S_f_) – to evaluate the impact of parasite infection on extra-pair paternity. The percentage of genetic offspring sired by the social father was compared to the total number of nestlings in the nest by dividing the number of genetic offspring by the total amount of offspring (G_o_/S_f_). White male paternity might be underestimated due to their promiscuous reproductive behavior (Grunst et al. (2018), potentially fathering offspring in a nest not included in this study.

### Parasite study

To determine infection status, a part of the mitochondrial cytochrome *b* of *Haemoproteus, Plasmodium*, and *Leucocytozoon* parasites was amplified with a nested PCR (Hellgren et al. 2004). All the malaria-positive samples were sequenced with Sanger sequencing (Hellgren et al. 2004). Finally, to identify the parasite genera, the sequences were compared in the existing parasites sequences database with the BLAST function of the MalAvi database (Bensch et al. 2009).

Parasitemia was evaluated using the general primers amplifying all three genera (*Plasmodium, Haemoproteus*, and *Leucocytozoon*) (Bell et al. 2015). First, all the samples were diluted to a similar DNA concentration (20 ng/µl), then the parasite DNA concentration was extrapolated based on a standard curve (1:100 to 1:100000) of a synthetic double-stranded DNA product of *Plasmodium relictum* as a positive control (Bell et al. 2015). Finally, all the reactions were performed in triplicate and then averaged the parasite concentration for each sample (see further details in Supporting information).

### Statistical analyses

All statistical analyses were performed with R software version V 4.1.2 (R Core Team 2021). We evaluated the impact of morph, sex, pair type, infection status, parasitemia, and population density (explanatory variables) on individual body condition (response variable) using a General Linear Mixed Model (GLMM) with a Gaussian distribution and identity link function. Parasitemia was a continuous predictor transformed as 1/-log (parasite concentration) to avoid very small numbers and potential convergence issues. Infection status was a categorical variable with two levels: single infected by *Plasmodium* or co-infected by *Plasmodium* and *Leucocytozoon*. Parasitemia and infection status were tested in separated models. Morph (tan or white), sex (male or female), and pair type (W×T or T×W) were also categorical variables with two levels. Population density was a continuous variable calculated for every year and habitat as the number of pairs per Ha. Year was included as a random factor.

To assess the effect of parasite infection on individual fitness, we also used GLMM for the next responses variables: number of fledglings (Poisson distribution, log link function), EPP (Binomial distribution, log link function), G_o_/S_f_ (Binomial distribution, log link function), nestling growth rate (Gaussian distribution, identity link function), and mean nestling body condition (Gaussian distribution and identity link function). Models were run separately for females and males to maximize sample size. All these models included as fixed predictors the parental variables: morph, body condition, infection status, and parasitemia. Clutch order (first or second) was also included as a fixed variable to account for potential differences between early- and late-season breeding attempts. We initially included the variable year as a random factor in all models, but removed it in most cases because year did not show variance. Missing information for individuals leads to variation in sample size available for each model.

Due to a large number of potential explanatory variables in our models, we used an information-theoretical approach based on the Akaike’s Information Criterion (AIC) to explore all possible combinations of explanatory variables and select the best models (Burnham and Anderson 2002) using the R package MuMIn version 1.43.17 (Bartoń 2020). Models were ranked according to their AIC, and we selected those with ΔAIC <4. We calculated the Akaike weight (ω) for all explanatory variables included in this subset of models. We selected those predictors with a medium or high effect (i.e., ω > 0.4) and ran a model containing only them for estimating their effects. The explanatory capacity of the latter model was calculated by its adjusted R^2^.

## RESULTS

Out of 90 adults we collected blood samples from, seven were infected with *Haemoproteus* (7.8%), 32 were infected with *Leucocytozoon* (35.6%), 74 were infected with *Plasmodium* (82.2%), and one solely with *Leucocytozoon* (1.1%). Since so few of our samples were non-infected (8/90), infected with *Haemoproteus* or solely with *Leucocytozoon*, we focused on samples infected only with *Plasmodium* (62.2%: 46/74) and co-infected with *Plasmodium*– *Leucocytozoon* (37.8%: 28/74). The average parasitemia level was 2.23×10^−8^ng/µl (SD=3.88×10^−8^; range=2.37×10^−10^ -1.88×10^−7^).

Based on the 74 adults selected, we collected 286 nestling blood samples from 88 nests over 17 years (Table S1; Supporting information). Sixty-nine nests were from the first clutch and 19 from the second clutch. We assigned the paternity of 268 nestlings genotyped, 49 of them being extra-pair offspring in 24 nests. Of the extra-pair offspring, 44 were assigned a genetic father, five of them were not allocated, their father being potentially an unknown individual from outside our study area.

### Body Condition

We found that individuals with a higher level of parasitemia show better body condition (Fig. 1), while co-infection has no relevant effects on body condition (Table 1). Furthermore, males always have better body conditions than females, while individuals breeding at higher population density are apparently in worse condition (the support for this variable was moderate). The selected explanatory variables in the models explain between 14 and 18% of body condition variability (Table 1).

**Table 1:** GLMM of the effect of parasite infection on adult body condition. SE is the standard error, and ω is the Akaike weight.

| | Predictors | Estimate ± SE | $\omega$ | Adjusted R<sup>2</sup> |
| --- | --- | --- | --- | --- |
| **Co-infection**<br>N=71 | Sex (male) | 1.03±0.27 | 0.991 | 0.175 |
| | Population density | -1.75±1.36 | 0.668 | |
| | Co-infection | 0.54±0.28 | 0.543 | |
| **Parasitemia**<br>N=71 | Sex (male) | 1.00±0.28 | 0.987 | 0.136 |
| | Parasitemia | 11.49±13.88 | 0.934 | |
| | Population density | -1.70±1.40 | 0.667 | |

**Figure 1:**
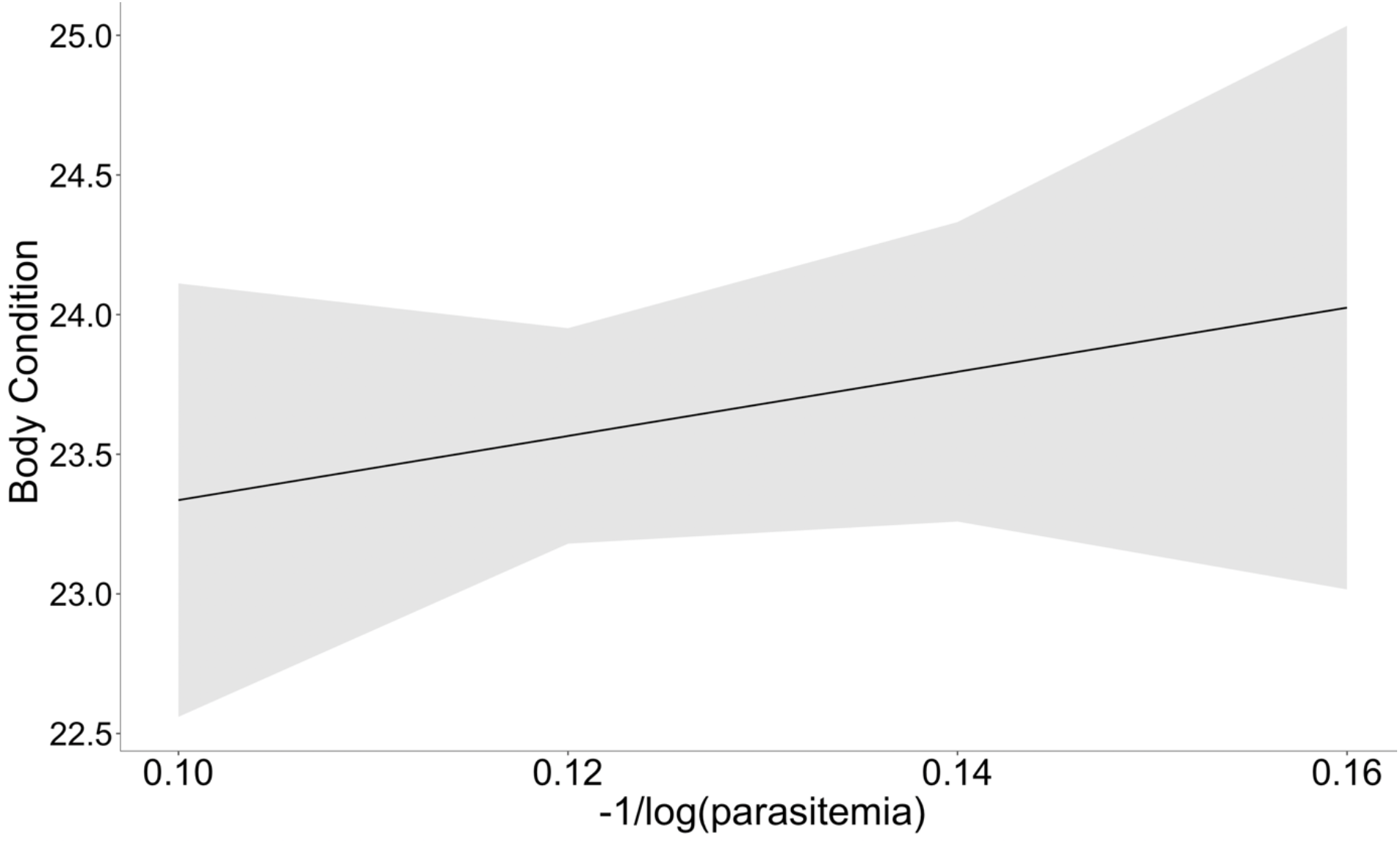
Effect of parasitemia on adult body condition. The shaded area represents the 95% confidence interval.

### Reproductive success

#### Fledglings per nest

Mother co-infection and parasitemia do not have any noticeable effect on the number of fledglings, as they have a weight too low to be included in the best models (Table S3a, Supporting information). White females and those in worse body conditions tend to raise more fledglings, but their effects are negligible and just explain 10% of the variation in breeding success of females (Table 2).

**Table 2:**
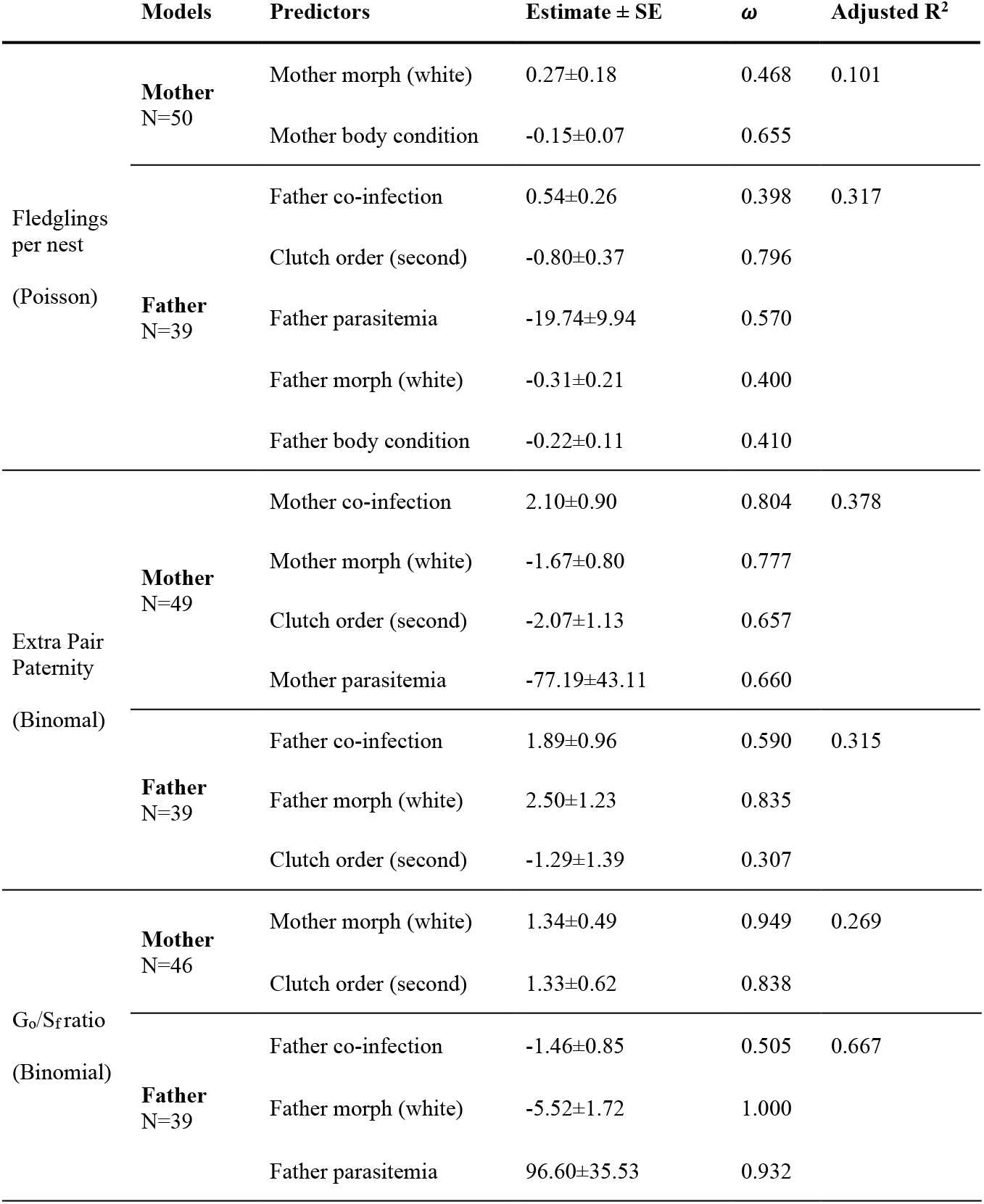
GLMM models of the effect of parasite infection on reproductive success. SE is the standard error, and ω is the Akaike weight.

Fathers’ co-infection and parasitemia are included in some of the best models for fledgling success, but both variables show a modest effect: co-infected males tend to have more fledglings, while those with a higher parasite load have less (Table 2). In the father’s models, we found that first clutches have more success than the next breeding attempt (Fig. 2). As in the case of females, father morph and body condition show negligible effects on the number of fledglings (Table 2).

**Figure 2:**
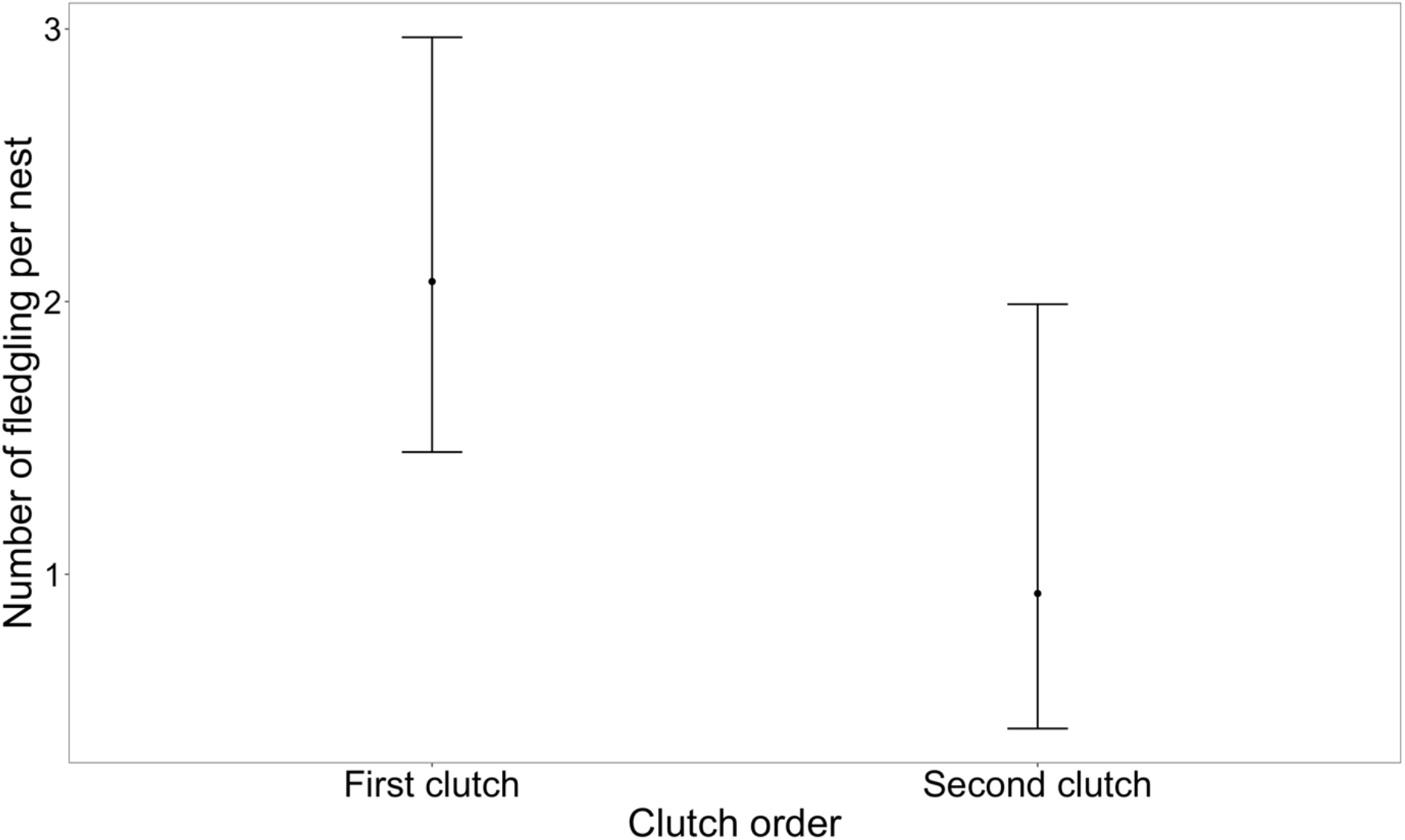
Average number of fledglings per nest in first and second clutches. Error bars denote the 95% confidence interval.

#### EPP per nest

Mothers’ co-infection have the most substantial effect on the probability of EPP (Table 2). Co-infected and tan females have more siblings from different males than their social mates (Fig. 3). However, higher levels of parasitemia and second clutches are related to a smaller chance of EPP. Overall, a notable 38% of the variability in the presence of EPP is explained in the mothers’ model (Table 2).

**Figure 3:**
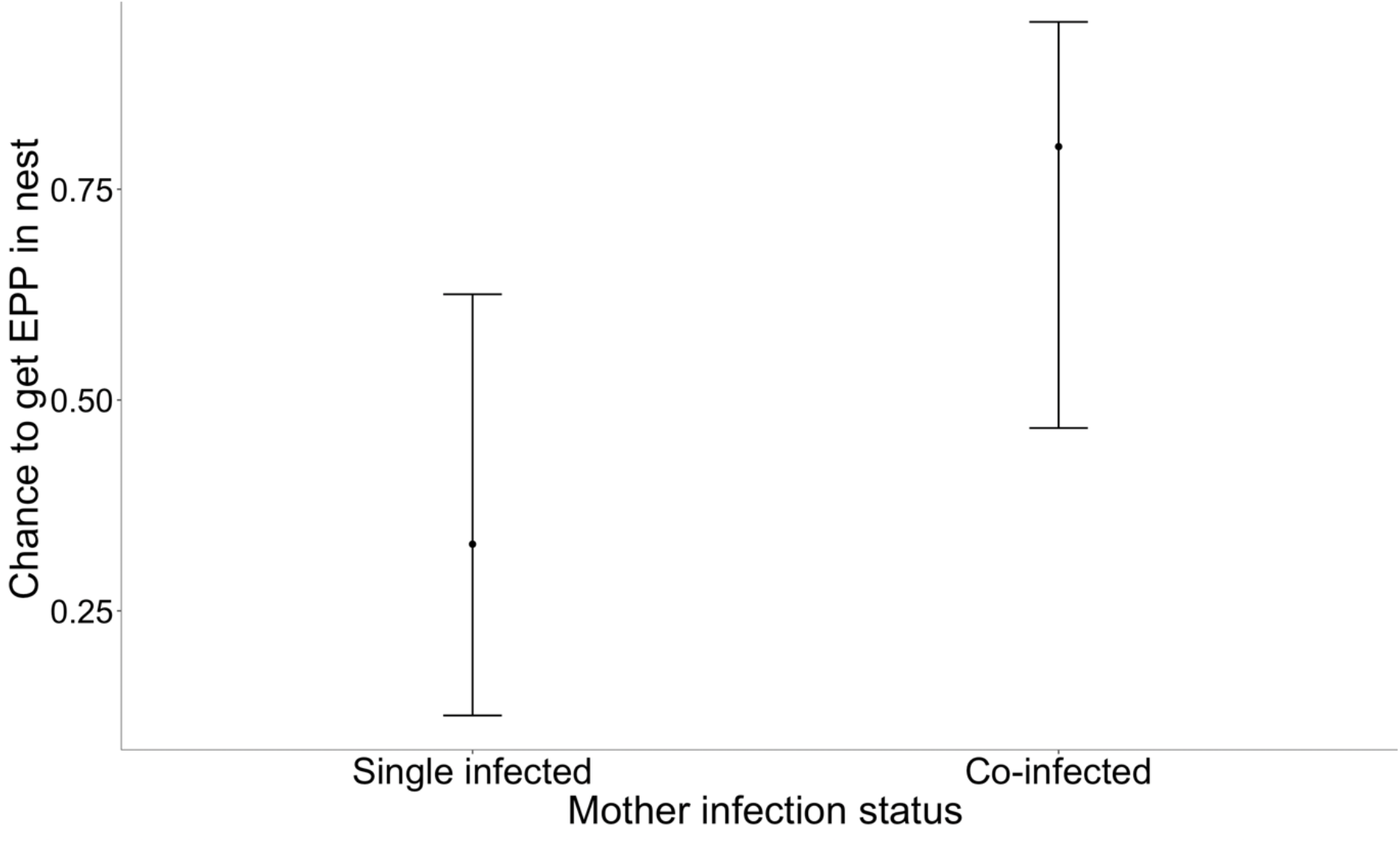
Average chance of extra-pair paternity occurrence in the nest for females with a single infection by *Plasmodium* or double infection by *Plasmodium* and *Leucocytozoon*. Error bars denote the 95% confidence interval.

Co-infected males also show more chances to suffer from EPP, but the effect of this variable is less relevant than in the females’ model (Table 2). Similarly, parasitemia does not show any relevant impact on EPP (Table S4b, Supporting information). The most pertinent factor for males was their morph, with white males having higher chances of raising illegitimate chicks (Table 2).

#### Ratio of genetic offspring belonging to the social father (G_o_/S_f_)

Mother’s parasites do not have any noticeable effect on the G_o_/S_f_ ratio (Table S5a, Supporting information). Mother morph and clutch order are the relevant predictors for G_o_/S_f_. White females and second clutches show the highest ratios (Table 2).

However, the father’s parasites play a prominent role in the G_o_/S_f_ ratio. Parasitemia show a marked positive effect, while co-infection has a medium adverse effect (Table 2). Thus, a higher parasite load increases the G_o_/S_f_ ratio (Fig. 4), while co-infections reduce it. Father’s morph is also essential for this model, as white males show a markedly lower G_o_/S_f_ ratio than tan males. Overall, the father’s model explains a noteworthy 67% of G_o_/S_f_ ratio variability among nests (Table 2).

**Figure 4:**
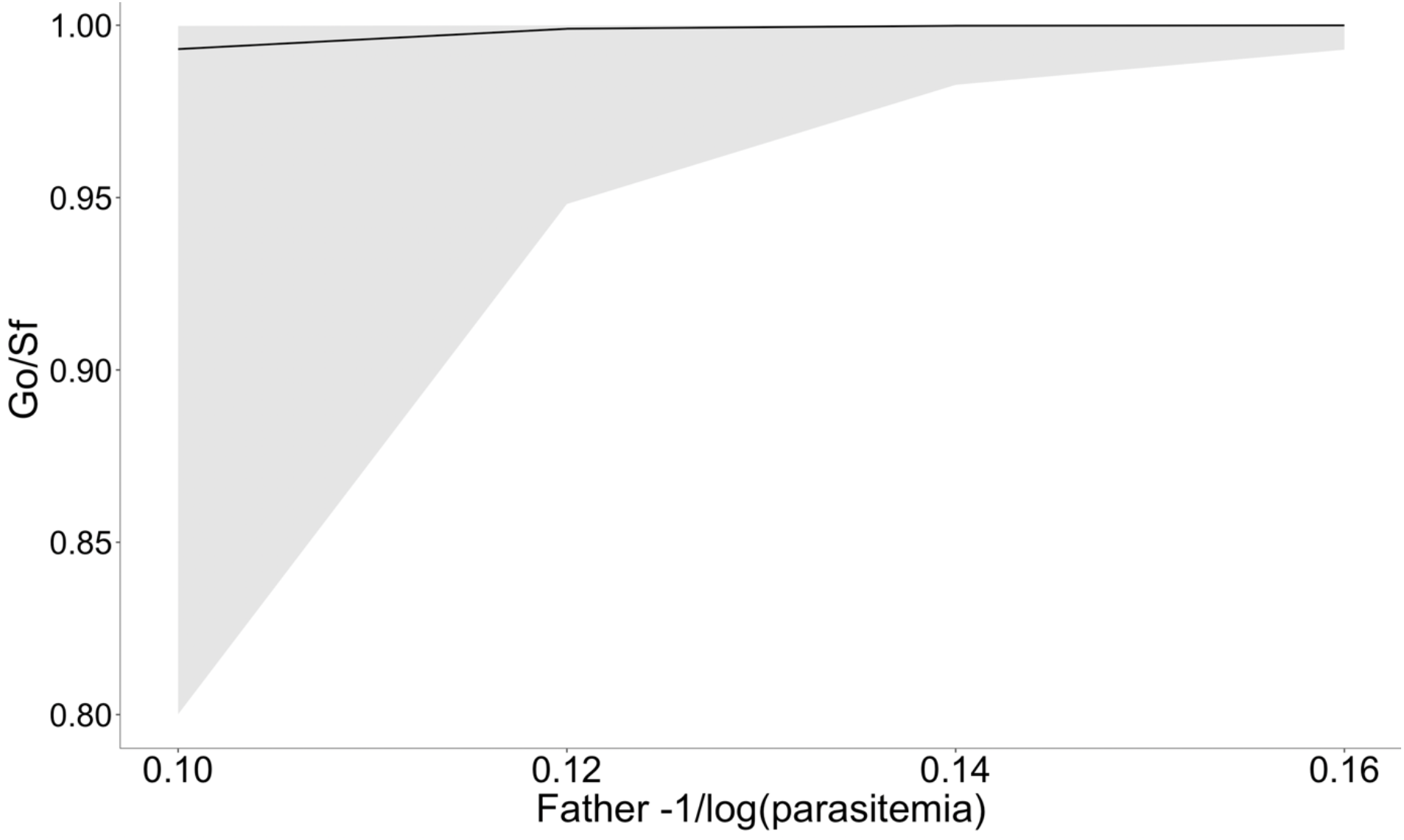
Effect of father parasitemia on the proportion of genetic offspring in nestlings (G_o_/S_f_). The shaded area represents the 95% confidence interval.

### Effect on nestling condition

#### Growth rate

Mothers’ parasite infection plays a role in the mean nestling growth rate variation. Co-infected mothers have nestlings with a higher mean growth rate, but the impact of co-infection is modest. In contrast, mother parasitemia has the most substantial effect, with mothers with high parasitemia having nestlings with higher growth rates (Fig. 5a). In this model, clutch order plays a medium role where first clutches show higher growth rates than the second ones (Table 3).

**Table 3:**
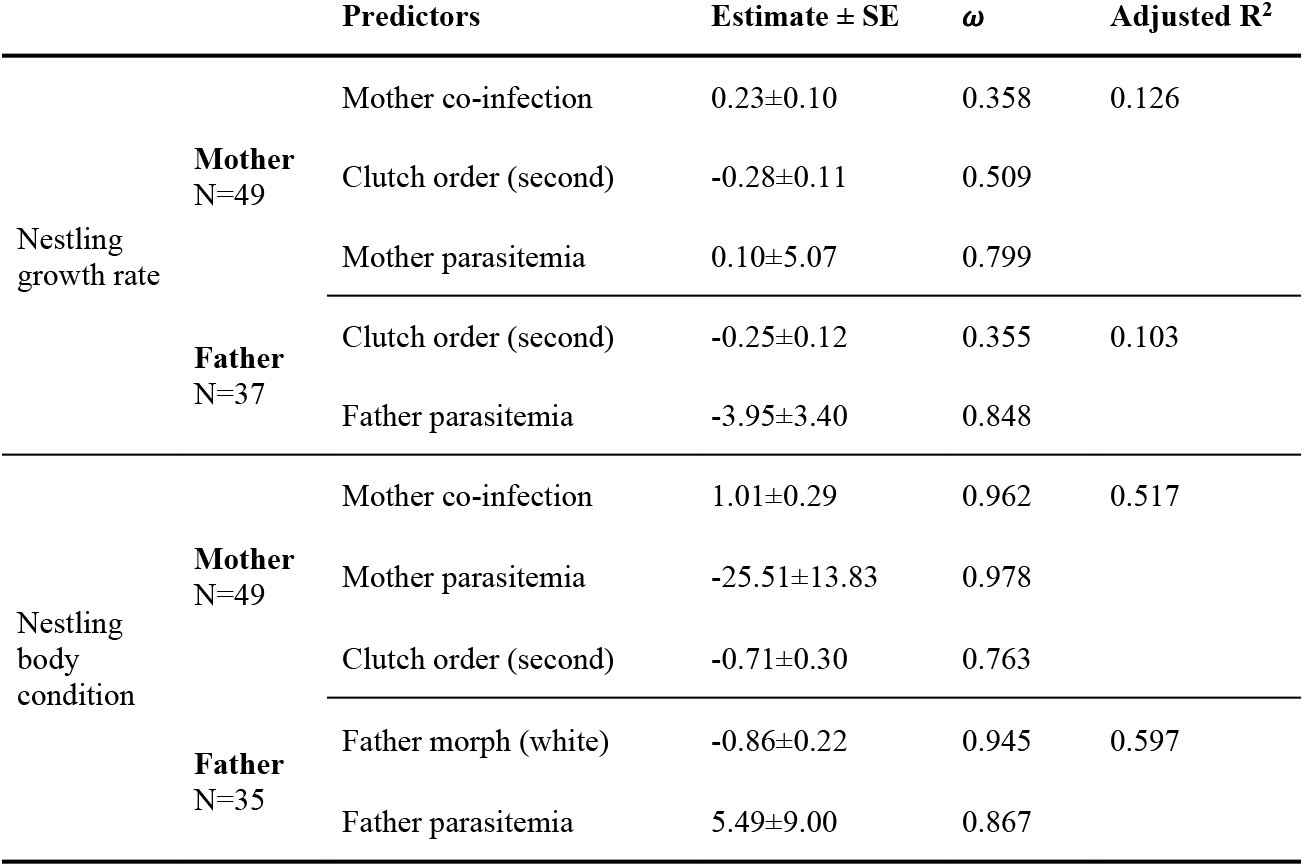
GLMM of the effect of parental infection on nestling growth rate and nestling body condition. SE is the standard error, and ω is the Akaike weight.

| | | Predictors | Estimate ± SE | $\omega$ | Adjusted R<sup>2</sup> |
| --- | --- | --- | --- | --- | --- |
| Nestling<br>growth rate | **Mother**<br>N=49 | Mother co-infection | 0.23±0.10 | 0.358 | 0.126 |
| | | Clutch order (second) | -0.28±0.11 | 0.509 | |
| | | Mother parasitemia | 0.10±5.07 | 0.799 | |
| | **Father**<br>N=37 | Clutch order (second) | -0.25±0.12 | 0.355 | 0.103 |
| | | Father parasitemia | -3.95±3.40 | 0.848 | |
| Nestling<br>body<br>condition | **Mother**<br>N=49 | Mother co-infection | 1.01±0.29 | 0.962 | 0.517 |
| | | Mother parasitemia | -25.51±13.83 | 0.978 | |
| | | Clutch order (second) | -0.71±0.30 | 0.763 | |
| | **Father**<br>N=35 | Father morph (white) | -0.86±0.22 | 0.945 | 0.597 |
| | | Father parasitemia | 5.49±9.00 | 0.867 | |

**Figure 5:**
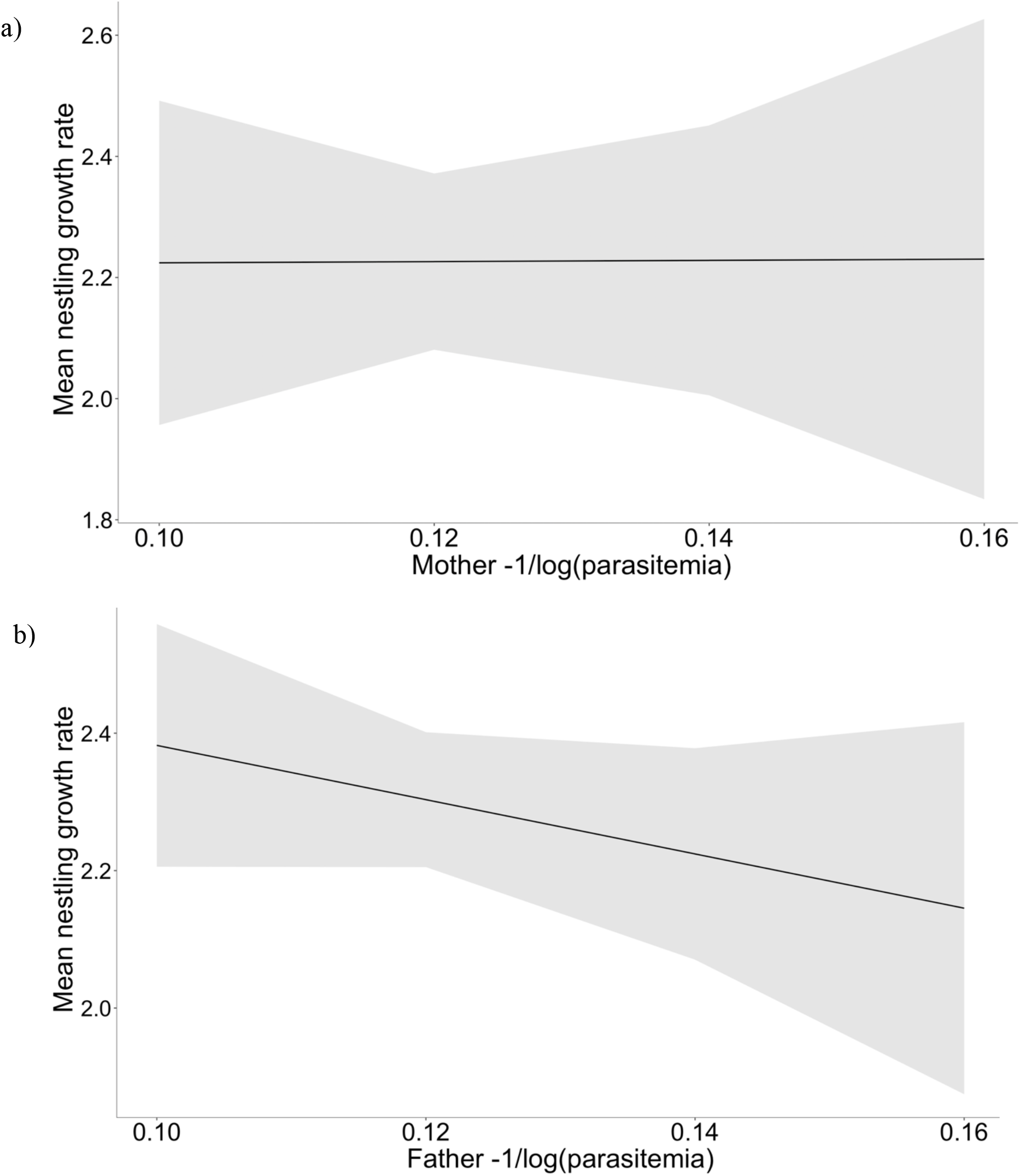
Effect of mother (a) and father (b) parasitemia on their nestlings’ growth rate. Shaded areas represent the 95% confidence interval.

Fathers’ co-infection status does not have any noticeable effect on nestling growth (Table S6b, Supporting information), but father parasitemia has a strong effect. Fathers with high parasitemia have a mean nestling growth rate lower than fathers with reduced parasitemia (Fig. 5b). Clutch order has similar effects to those previously seen in the mother model, but only with a very modest effect (Table 3).

Overall, both models explain between 10% to 13% of the variation in growth rate in nestlings (Table 3).

#### Body condition

Mothers’ infection plays a prominent role in the variation of nestling body condition. Co-infected mothers have nestlings with better body condition (Fig. 6a). On contrary, nestlings cared for by females with high parasitemia and from second clutches have lower body condition (Table 3, Fig. 6b).

**Figure 6:**
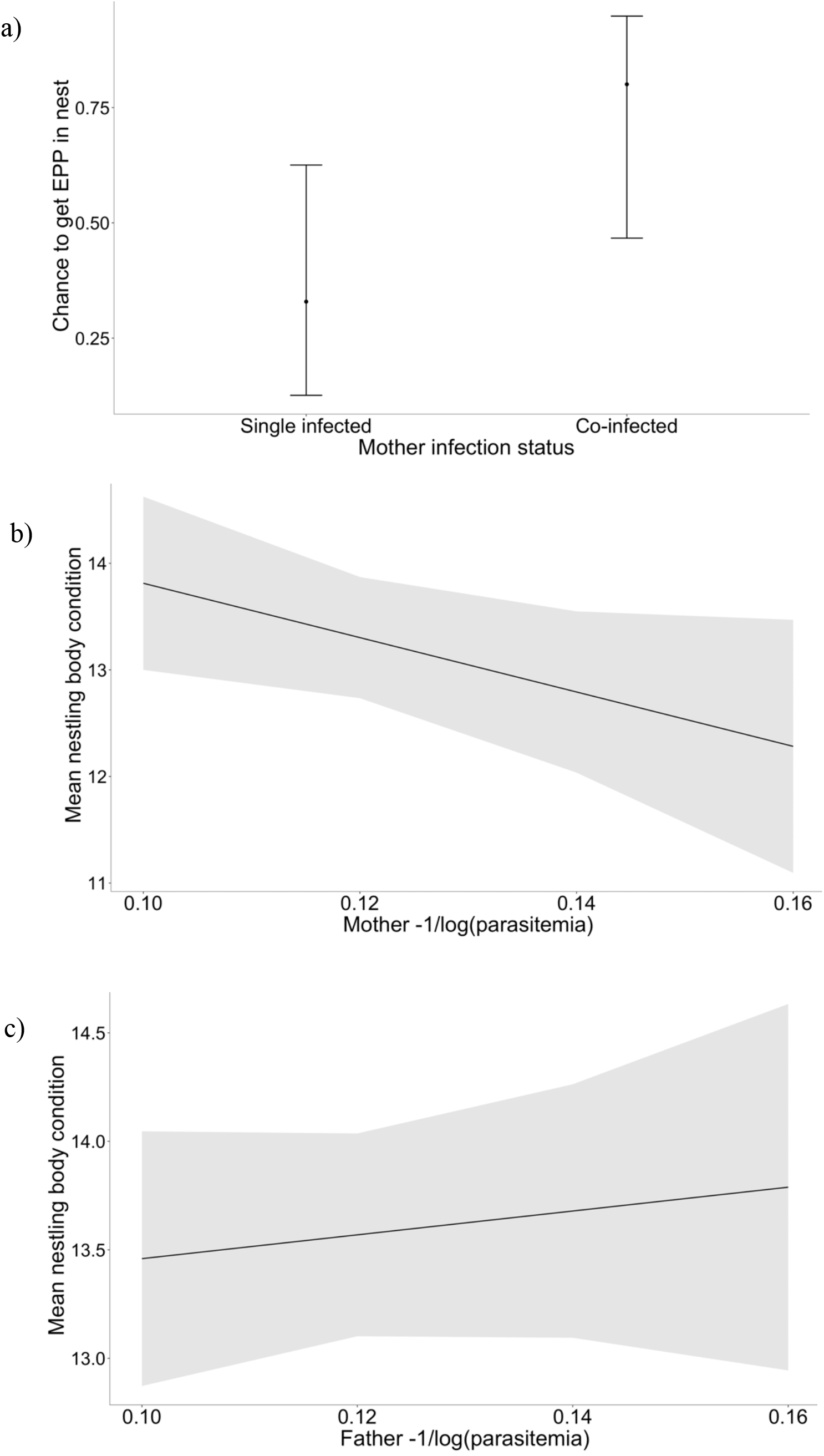
Effect of mother infection status (a), mother parasitemia (b), and father parasitemia (c) on the nestling body condition. Error bars and shaded area represent the 95% confidence interval.

Fathers’ co-infection status has a too low weight to be included in the best model, not having any noticeable effect on the nestling body condition (Table S7b, Supporting information). However, fathers’ parasitemia and morph have a substantial impact on nestling body condition. Fathers with high parasitemia and tan morph have nestlings in better condition than the rest of the males (Table 3, Fig. 6c). Altogether, the mothers’ model explains 52%, and the fathers’ model explains 60% of the variation in body condition in nests (Table 3).

## DISCUSSION

In this study, we investigated how haemosporidian parasite infection influences host body condition and reproductive performance in terms of breeding success, quality of the offspring and extra-pair paternity in white-throated sparrow. We found that co-infection by *Plasmodium* and *Leucocytozoon* positively impacts host reproductive success by increasing extra-pair paternity (EPP) and nestling quality, suggesting a switch in reproductive strategy to maximize the current reproduction, supporting the terminal investment hypothesis (Part et al. 1992). On the other hand, high levels of parasites in the blood (i.e., parasitemia) impact reproductive success by increasing the actual percentage of genetic offspring (the G_o_/S_f_ ratio), but have mixed effects on nestling quality.

### Parasite infection and host body condition

Contrary to our expectations, co-infection status does not influence body condition in adult white-throated sparrows. Our results do not follow the observation made in other bird species for single and co-infection. In tawny pipits (*Anthus campestris*), Calero-Riestra and García (2016) found that infection by *Haemoproteus* or *Plasmodium* negatively influence body condition. When co-infected with *Haemoproteus* and *Plasmodium*, house martins (*Delichon urbicum*) show impaired body condition (Marzal et al. 2008). But when tested in North American birds, including white-throated sparrow, Astudillo et al. (2013) observed that neither *Plasmodium* nor *Leucocytozoon* prevalence affect the body mass index of the sampled birds. However, the body condition was measured differently in each of those studies, and the contrasting results might be caused by other parasite infections having a contrasting effect on host body condition (Atkinson and van Riper III 1991, Bentz et al. 2006).

While co-infection does not impact body condition of white-throated sparrows, parasitemia has a positive effect as individuals with high parasitemia have better body condition. Higher parasitemia (*Plasmodium* or *Haemoproteus*) has been shown in white-necked thrush (*Turdus albicollis*) and in blue tits (*Cyanistes caeruleus*) to have a negative influence on body condition (Merino et al. 2000, De La Torre et al. 2020). Our contrasting result between high parasitemia and high body condition could be due to the cost of immunity. Individuals with low body condition and low parasitemia might pay the price of immune response stimulation, expected to be highly energy-consuming. In tawny owl, grey individuals have a higher immune response than rufous ones with the same level of *Leucocytozoon* parasitemia (Galeotti and Sacchi 2003). This higher immune response leaves the grey morph with decreased parasitemia and a body condition negatively associated with increased eosinophil concentration, leaving the brown morph with higher parasitemia (Galeotti and Sacchi 2003). Similarly, the activation of the immune system to suppress infection is associated with a loss of body condition in grey females with nestlings, while it does not impact brown females body condition (Karell et al. 2011). These differences in our findings and those of De La Torre et al. (2020) and Merino et al. (2000) could be caused by the combination of the parasitemia of *Plasmodium* and *Leucocytozoon*, as our test does not discriminate between parasite genera, since parasitemia of each genus could vary widely causing different responses by the host.

### Parasite infection and reproductive success

While parental co-infection status does not influence the number of fledglings per nest and the ratio of genetic offspring belonging to the social father, mothers’ co-infection status increases the chance of having an EPP in their nests. Co-infection could stimulate female motivation to seek extra-pair copulation to increase the genetic diversity of their progeny and diversify their defenses against multiple parasite infections, following the “good genes” hypothesis (Podmokła et al. 2015, Hamilton and Zuk 1982, Forstmeier et al. 2014). In males, we could not confirm this hypothesis because we did not have the blood samples to evaluate the parasite infection status of the external father, although we identified most of the fathers of extra-pair paternity offspring.

Males with high parasitemia have a higher ratio of genetic offspring belonging to the social father. High parasitemia could reduce the mobility of parasitized males (Mukhin et al. 2016), through severe costs of parasite multiplication, leading to the destruction of red blood cells and increased anemia (Schoenle et al. 2017). In that case, males with reduced mobility would be less prone to abandon their territories, and indirectly they can invest more in females guarding and preserving their paternity. Thus, males could be induced by parasites to switch their reproductive strategies. Moreover, heavily infected males that would not survive the following season should be favored by selection if performing terminal reproductive investments and increasing their immediate fitness (Schwanz 2008).

### Parental infection and offspring

Parental infections have a substantial impact on white-throated sparrow nestling quality. We found that co-infected females have nestlings in better body condition than single infected ones. Previous research suggested that when *Plasmodium* or *Haemoproteus* infect both parents, they sire larger and heavier nestlings than when only one or none of the parents are infected (Podmokła et al. 2014). However, when parental co-infection was tested on nestling body mass, it shows no effect, only their body condition affects nestling body condition (Pigeault et al. 2020). The increase in nestling body condition in co-infected females might be linked to a switch in reproductive strategy where highly parasitized females perform a terminal investment, optimizing the survival of the current brood (Kulma et al. 2014, Schwanz 2008).

Parental parasitemia influences nestling quality differently. While mothers with high parasitemia influence nestling body condition negatively but nestling growth rate positively, fathers with high parasitemia have an opposite effect. However, the mothers’ positive influence on nestling growth rate is low. Highly infected females exhausted by nest incubation might have a significant immune reaction to parasite infection, decreasing their investment and food nestling provisioning, causing lower nestling body condition (Grzędzicka 2017). In blue tit, females treated with antimalarial drugs have their parasitemia reduced, which results in lower brood weight inequalities (Knowles et al. 2010). Parental provision rate is expected to decrease with infection and high parasitemia, if parasites impact parental performance (Buchanan et al. 1999, Knowles et al. 2010). On the other hand, fathers with high parasitemia have nestlings with low growth rates and increased body condition. With lower chances of surviving due to high parasitemia, males might adopt the high energetic task of defending their territory and invest more in food provisioning, leading to heavier offspring.

### Effect of other variables on reproductive success

White-throated sparrows present two different reproductive strategies as a result of their disassortative mating. Pairs formed by tan males and white females (T×W) provide better parental care than pairs of white males and tan females (W×T). In the latter, females primarily attend to the offspring, while in the former both parents share parental duties (Tuttle 2003). White males spend more time in territory defense and seeking extra-pair copulation, effectively leaving tan females as “single mothers” (Grunst et al. 2018). These two alternative strategies are reflected in our observation of morph effect on reproductive success and nestling quality. W×T pairs have higher extra-pair paternity and a lower ratio of genetic offspring belonging to the social father with lower body condition nestlings than T×W pairs. Our results are congruent with the pair-type reproductive strategies of the white-throated sparrow, where W×T pairs are more promiscuous, while T×W pairs have a more monogamous relationship involving mate guarding (Grunst et al. 2019b).

We also observed that second clutches have fewer successful fledglings and a higher ratio of genetic offspring belonging to the social father than first clutches. First clutches are quite synchronous with every pair building and laying eggs approximately at the same time. As the breeding season progresses, subsequent clutches become more asynchronous. Females that fail their first clutch are busy building a new nest, while successfully reproducing males might be less keen on seeking extra-pair copulation because they take care of their first clutch offspring. Additionally, second clutches yield nestlings with a lower body condition. Decreased body condition could be due to the potential decrease of food availability later in the season or parental exhaustion after raising the first clutch, making a worse food provisioning (Kulma et al. 2014).

## CONCLUSION

We show that regardless of morph, co-infected white-throated sparrows increase their immediate reproductive success by increasing the mean nestling growth rate and body condition, as well as extra-pair paternity. This increase in reproductive success with the addition of genetic diversity due to EPP might be a mechanism to cope with the cost of multiple parasite infections, switching to a terminal investment reproductive strategy (Agnew et al. 2000, Kulma et al. 2014). Previous studies suggesting this switch in reproductive strategies mainly compared infected with co-infected individuals (Norte et al. 2009, Pigeault et al. 2020). Few investigated the differences in reproductive strategies between uninfected, single infected, and co-infected (Marzal et al. 2008). Host survival challenged by co-infection might trigger terminal investments to maximize their current reproductive output (Marzal et al. 2008). However, the effect of parasitemia on reproductive success follows the expectation that high levels of the parasite in blood would increase the ratio of genetic offspring belonging to the social father and have a contrasting impact on nestling quality. The contrasting impact of infection on nestling quality between mothers and fathers could result from female higher energetic input throughout the breeding season, building nest, laying, and incubating eggs. At the same time, males invest energy only when brood feeding.

## DATA AVAILABILITY STATEMENT

The data associated with this research are openly available in Dryad at DOI.. upon publication

## CONFLICT OF INTEREST

The authors have no conflict of interest to declare.

## ETHICS STATEMENT

This research followed the laws of New York state, the state of Indiana, and the US federal government (USGS Fish & Wildlife Federal Banding Permit #24105 to RA Gonser). This research was conducted under the approval of Indiana State University’s Institutional Animal Care and Use Committee (Protocol #56218-1).

## FUNDING STATEMENT

Funding was provided by the Indiana Academy of Science, college of graduate and professional studies, and Center for Genomic Advocacy at Indiana State University (to ZD and RAG) and by National Science Foundation grant DUE-0934648 (to Elaina M. Tuttle).

## AUTHOR’S CONTRIBUTIONS

ZD, HFG, and RAG conceived the ideas and designed the methodology; ZD and KRS collected the data; ZD and OG analyzed the data; ZD led the writing of the manuscript. All authors contributed critically to the drafts and gave final approval for publication.

## ACKNOWLEDGEMENTS

We would like to thank Elaina M. Tuttle for her lifetime engagement in the white-throated sparrow project and the Tuttle-Gonser lab members from 2002-2021 for their field and laboratory work.

## Notes

https://doi.org/10.6084/m9.figshare.19145951

### Competing Interest Statement

The authors have declared no competing interest.

## REFERENCES

Agnew, P., Koella, J. C. and Michalakis, Y. 2000. Host life history responses to parasitism. - Microb. Infect., 2: 891–896. https://doi.org/10.1016/S1286-4579(00)00389-0

Astudillo, V. G., Hernández, S. M., Kistler, W. M., Boone, S. L., Lipp, E. K., Shrestha, S. and Yabsley, M. J. 2013. Spatial, temporal, molecular, and intraspecific differences of haemoparasite infection and relevant selected physiological parameters of wild birds in Georgia, USA. - Int. J. Parasitol.: Parasit. Wildl., 2: 178–189. https://doi.org/10.1016/j.ijppaw.2013.04.005

Atkinson, C. T. and Van Riper Iii, C. 1991. Pathogenicity and epizootiology of avian haematozoa: Plasmodium, Haemoproteus, and Leucocytozoon. Bird-parasite interactions: Ecology, evolution, and behavior. Oxford University Press, London, pp. 19–48.

Badás, E. P., Autor, A., Martínez, J., Rivero-De Aguilar, J. and Merino, S. 2020. Individual Quality and Extra-Pair Paternity in the Blue Tit: Sexy Males Bear the Costs. - Evolution, 74: 559–572. https://doi.org/10.1111/evo.13925

Bartoń, K. 2020. MuMIn: Multi-Model Inference. https://cran.r-project.org/web/packages/MuMIn/index.html

Bell, J. A., Weckstein, J. D., Fecchio, A. and Tkach, V. V. 2015. A new real-time PCR protocol for detection of avian haemosporidians. - Parasites Vectors, 8: 9. https://doi.org/10.1186/s13071-015-0993-0

Bensch, S., Hellgren, O. and Pérez-Tris, J. 2009. MalAvi: a public database of malaria parasites and related haemosporidians in avian hosts based on mitochondrial cytochrome b lineages. - Mol. Ecol. Resour., 9: 1353–1358. https://doi.org/10.1111/j.1755-0998.2009.02692.x

Bentz, S., Rigaud, T., Barroca, M., Martin-Laurent, F., Bru, D., Moreau, J. and Faivre, B. 2006. Sensitive measure of prevalence and parasitaemia of haemosporidia from European blackbird (Turdus merula) populations: value of PCR-RFLP and quantitative PCR. - Parasitology, 133: 685–692. https://doi.org/10.1017/s0031182006001090

Bose, J., Kloesener, M. H. and Schulte, R. D. 2016. Multiple-genotype infections and their complex effect on virulence. - Zoology, 119: 339–349. https://doi.org/10.1016/j.zool.2016.06.003

Boyd, R., Kelly, T., Macdougall-Shackleton, S. and Macdougall-Shackleton, E. 2018. Alternative reproductive strategies in white-throated sparrows are associated with differences in parasite load following experimental infection. - Biol. Lett., 14: 20180194. https://doi.org/10.1098/rsbl.2018.0194

Buchanan, K., Catchpole, C., Lewis, J. and Lodge, A. 1999. Song as an indicator of parasitism in the sedge warbler. - Anim. Behav., 57: 307–314. https://doi.org/10.1006/anbe.1998.0969

Burnham, K. P. and Anderson, D. R. 2002. Model selection and multimodel inference: A Practical Information-Theoretic Approach, Springer, New York, NY.

Calero-Riestra, M. and García, J. T. 2016. Sex-dependent differences in avian malaria prevalence and consequences of infections on nestling growth and adult condition in the Tawny pipit, Anthus campestris. - Malar. J., 15: 178. https://doi.org/10.1186/s12936-016-1220-y

Cuevas, E., Orellana-Penailillo, C., Botero-Delgadillo, E., Espindola-Hernandez, P., Vasquez, R. A. and Quirici, V. 2021. Influence of the haemosporidian Leucocytozoon spp. over reproductive output in a wild Neotropical passerine, the Thorn-tailed Rayadito (Aphrastura spinicauda). - Ibis, 163: 948–961. https://doi.org/10.1111/ibi.12934

De La Torre, G. M., Freitas, F. F., Fratoni, R. D. O., Guaraldo, A. D. C., Dutra, D. D. A., Braga, M. and Manica, L. T. 2020. Hemoparasites and their relation to body condition and plumage coloration of the White-necked thrush (Turdus albicollis). - Ethol. Ecol. Evol., 32: 509–526. https://doi.org/10.1080/03949370.2020.1769739

Fallon, S. M., Ricklefs, R. E., Swanson, B. and Bermingham, E. 2003. Detecting avian malaria: an improved polymerase chain reaction diagnostic. - J. Parasitol., 89: 1044–1047. https://doi.org/10.1645/GE-3157

Figuerola, J., Munoz, E., Gutierrez, R. and Ferrer, D. 1999. Blood Parasites, Leucocytes and Plumage Brightness in the Cirl Bunting, Emberiza cirlus. - Funct. Ecol., 13: 594–601. http://www.jstor.org/stable/2656310

Forstmeier, W., Nakagawa, S., Griffith, S. C. and Kempenaers, B. 2014. Female extra-pair mating: adaptation or genetic constraint? - Trends Ecol. Evol., 29: 456–464. https://doi.org/10.1016/j.tree.2014.05.005

Galeotti, P. and Sacchi, R. 2003. Differential parasitaemia in the tawny owl (Strix aluco): effects of colour morph and habitat. - J. Zool., 261: 91–99. https://doi.org/10.1017/s0952836903003960

Gangoso, L., Gutiérrez-López, R., Puente, J. M.-d. l. and Figuerola, J. 2016. Genetic colour polymorphism is associated with avian malarial infections. - Biol. Lett., 12: 20160839. https://doi.org/10.1098/rsbl.2016.0839

Gilman, S., Blumstein, D. T. and Foufopoulos, J. 2007. The Effect of Hemosporidian Infections on White-Crowned Sparrow Singing Behavior. - Ethology, 113: 437–445. https://doi.org/10.1111/j.1439-0310.2006.01341.x

Grunst, A. S., Grunst, M. L., Formica, V. A., Korody, M. L., Betuel, A. M., Barcelo-Serra, M., Ford, S., Gonser, R. A. and Tuttle, E. M. 2018. Morph-Specific Patterns of Reproductive Senescence: Connections to Discrete Reproductive Strategies. - Am. Nat., 191: 744–755. https://doi.org/10.1086/697377

Grunst, A. S., Grunst, M. L., Gonser, R. A. and Tuttle, E. M. 2019a. Developmental stress and telomere dynamics in a genetically polymorphic species. - J. Evol. Biol., 32: 134–143. https://doi.org/10.1111/jeb.13400

Grunst, A. S., Grunst, M. L., Korody, M. L., Forrette, L. M., Gonser, R. A. and Tuttle, E. M. 2019b. Extrapair mating and the strength of sexual selection: insights from a polymorphic species. - Behav. Ecol., 30: 278–290. https://doi.org/10.1093/beheco/ary160

Grzędzicka, E. 2017. Immune challenge of female great tits at nests affects provisioning and body conditions of their offspring. - Acta Ethol, 20: 223–233. https://doi.org/10.1007/s10211-017-0265-4

Hamilton, W. D. and Zuk, M. 1982. Heritable true fitness and bright birds: a role for parasites? - Science, 218: 384–387. https://doi.org/10.1126/science.7123238

Hellgren, O., Waldenström, J.MJ. and Bensch, S. 2004. A new PCR assay for simultaneous studies of Leucocytozoon, Plasmodium, and Haemoproteus from avian blood. - J. Parasitol., 90: 797–802. https://doi.org/10.1645/ge-184r1

Ishtiaq, F., Rao, M., Huang, X. and Bensch, S. 2017. Estimating prevalence of avian haemosporidians in natural populations: a comparative study on screening protocols. - Parasites Vectors, 10: 10. https://doi.org/10.1186/s13071-017-2066-z

Jacobs, A. C., Fair, J. M. and Zuk, M. 2015. Parasite infection, but not immune response, influences paternity in western bluebirds. - Behav. Ecol. Sociobiol., 69: 193–203. https://doi.org/10.1007/s00265-014-1832-6

Karell, P., Ahola, K., Karstinen, T., Kolunen, H., Siitari, H. and Brommer, J. 2011. Blood parasites mediate morph-specific maintenance costs in a colour polymorphic wild bird. - J. Evol. Biol., 24: 1783–1792. https://doi.org/10.1111/j.1420-9101.2011.02308.x

Karell, P., Bensch, S., Ahola, K. and Asghar, M. 2017. Pale and dark morphs of tawny owls show different patterns of telomere dynamics in relation to disease status. - Proc. R. Soc. B: Biol. Sci., 284: 20171127. https://doi.org/10.1098/rspb.2017.1127

Karell, P., Pietiäinen, H., Siitari, H. and Brommer, J. 2007. A possible link between parasite defence and residual reproduction. - J. Evol. Biol., 20: 2248–2252. https://doi.org/10.1111/j.1420-9101.2007.01423.x

Knowles, S. C., Palinauskas, V. and Sheldon, B. 2010. Chronic malaria infections increase family inequalities and reduce parental fitness: experimental evidence from a wild bird population. - J. Evol. Biol., 23: 557–569. https://doi.org/10.1111/j.1420-9101.2009.01920.x

Knowles, S. C., Wood, M. J., Alves, R., Wilkin, T. A., Bensch, S. and Sheldon, B. C. 2011. Molecular epidemiology of malaria prevalence and parasitaemia in a wild bird population. - Mol. Ecol., 20: 1062–1076. https://doi.org/10.1111/j.1365-294X.2010.04909.x

Kulma, K., Low, M., Bensch, S. and Qvarnström, A. 2014. Malaria-Infected Female Collared Flycatchers (Ficedula albicollis) Do Not Pay the Cost of Late Breeding. - PLoS ONE, 9: e85822. https://doi.org/10.1371/journal.pone.0085822

Lapointe, D. A., Atkinson, C. T. and Samuel, M. D. 2012. Ecology and conservation biology of avian malaria. - Ann. N. Y. Acad. Sci., 1249: 211–226. https://doi.org/10.1111/j.1749-6632.2011.06431.x

Lei, B., Amar, A., Koeslag, A., Gous, T. A. and Tate, G. J. 2014. Differential Haemoparasite Intensity between Black Sparrowhawk (Accipiter melanoleucus) Morphs Suggests an Adaptive Function for Polymorphism. - PLoS ONE, 8: e81607. https://doi.org/10.1371/journal.pone.0081607

Longmire, J. L., Gee, G. F., Hardekopf, C. L. and Mark, G. A. 1992. Establishing Paternity in Whooping Cranes (Grus americana) by DNA Analysis. - Auk, 109: 522–529. https://doi.org/10.1093/auk/109.3.522

Lowther, J. K. 1962. Colour and Behavioural Polymorphism in the White-throated Sparrow, Zonotrichia Albicollis (Gmelin). University of Toronto.

Macdougall-Shackleton, E. A., Derryberry, E. P. and Hahn, T. P. 2002. Nonlocal male mountain white-crowned sparrows have lower paternity and higher parasite loads than males singing local dialect. - Behav. Ecol., 13: 682–689. https://doi.org/10.1093/beheco/13.5.682

Marzal, A., Bensch, S., Reviriego, M., Balbontin, J. and De Lope, F. 2008. Effects of malaria double infection in birds: one plus one is not two. - J. Evol. Biol., 21: 979–987. https://doi.org/10.1111/j.1420-9101.2008.01545.x

Marzal, A., De Lope, F., Navarro, C. and Møller, A. P. 2005. Malarial parasites decrease reproductive success: an experimental study in a passerine bird. - Oecologia, 142: 541–545. https://doi.org/10.1007/s00442-004-1757-2

Merino, S., Moreno, J., Sanz, J. J. and Arriero, E. 2000. Are avian blood parasites pathogenic in the wild? A medication experiment in blue tits (Parus caeruleus). - Proc. R. Soc. B: Biol. Sci., 267: 2507–2510. https://doi.org/10.1098/rspb.2000.1312

Michopoulos, V., Maney, D. L., Morehouse, C. B. and Thomas, J. W. 2007. A genotyping assay to determine plumage morph in the white-throated sparrow (Zonotrichia albicollis). - Auk, 124: 1330–1335. https://doi.org/10.1093/auk/124.4.1330

Mukhin, A., Palinauskas, V., Platonova, E., Kobylkov, D., Vakoliuk, I. and Valkiūnas, G. 2016. The strategy to survive primary malaria infection: an experimental study on behavioural changes in parasitized birds. - PLoS ONE, 11: e0159216. https://doi.org/10.1371/journal.pone.0159216

Nordling, D., Andersson, M., Zohari, S. and Lars, G. 1998. Reproductive effort reduces specific immune response and parasite resistance. - Proc. R. Soc. B: Biol. Sci., 265: 1291–1298. https://doi.org/10.1098/rspb.1998.0432

Norte, A. C., Araújo, P. M., Sampaio, H. L., Sousa, J. P. and Ramos, J. A. 2009. Haematozoa infections in a Great Tit Parus major population in Central Portugal: relationships with breeding effort and health. - Ibis, 151: 677–688. https://doi.org/10.1111/j.1474-919X.2009.00960.x

Palinauskas, V., Žiegytė, R., Iezhova, T. A., Ilgūnas, M., Bernotienė, R. and Valkiūnas, G. 2016. Description, molecular characterisation, diagnostics and life cycle of Plasmodium elongatum (lineage pERIRUB01), the virulent avian malaria parasite. - Int. J. Parasitol., 46: 697–707. https://doi.org/10.1016/j.ijpara.2016.05.005

Part, T., Gustafsson, L. and Moreno, J. 1992. “Terminal Investment” and a Sexual Conflict in the Collared Flycatcher (Ficedula albicollis). - Am. Nat., 140: 868–882. https://doi.org/10.1086/285445

Peig, J. and Green, A. J. 2009. New perspectives for estimating body condition from mass/length data: the scaled mass index as an alternative method. - Oikos, 118: 1883–1891. https://doi.org/10.1111/j.1600-0706.2009.17643.x

Pigeault, R., Cozzarolo, C.-S., Glaizot, O. and Christe, P. 2020. Effect of age, haemosporidian infection and body condition on pair composition and reproductive success in Great Tits Parus major. - Ibis, 162: 613–626. https://doi.org/10.1111/ibi.12774

Pigeault, R., Cozzarolo, C. S., Choquet, R., Strehler, M., Jenkins, T., Delhaye, J., Bovet, L., Wassef, J., Glaizot, O. and Christe, P. 2018. Haemosporidian infection and co-infection affect host survival and reproduction in wild populations of great tits. - Int. J. Parasitol., 48: 1079–1087. https://doi.org/10.1016/j.ijpara.2018.06.007

Podmokła, E., Dubiec, A., Arct, A., Drobniak, S. M., Gustafsson, L. and Cichoń, M. 2015. Malaria infection status predicts extra-pair paternity in the blue tit. - J. Avian Biol., 46: 303–306. https://doi.org/10.1111/jav.00599

Podmokła, E., Dubiec, A., Drobniak, S. M., Arct, A., Gustafsson, L. and Cichoń, M. 2014. Avian malaria is associated with increased reproductive investment in the blue tit. - J. Avian Biol., 45: 219–224. https://doi.org/10.1111/j.1600-048X.2013.00284.x

Poulin, R. 2011. Evolutionary Ecology of Parasites: (Second Edition), Princeton University Press.

R Core Team, C. 2021. R: a language and environment for statistical computing. - R Foundation for statistical computing, Vienna.

Reaney, L. T. and Knell, R. J. 2010. Immune activation but not male quality affects female current reproductive investment in a dung beetle. - Behav. Ecol., 21: 1367–1372. https://doi.org/10.1093/beheco/arq139

Reinoso-Pérez, M. T., Canales-Delgadillo, J. C., Chapa-Vargas, L. and Riego-Ruiz, L. 2016. Haemosporidian parasite prevalence, parasitemia, and diversity in three resident bird species at a shrubland dominated landscape of the Mexican highland plateau. - Parasites Vectors, 9: 12. https://doi.org/10.1186/s13071-016-1569-3

Reinoso-Pérez, M. T., Dhondt, K. V., Sydenstricker, A. V., Heylen, D. and Dhondt, A. A. 2020. Complex interactions between bacteria and haemosporidia in coinfected hosts: An experiment. - Ecol. Evol., 10: 5801–5814. https://doi.org/10.1002/ece3.6318

Roulin, A. 2004. The evolution, maintenance and adaptive function of genetic colour polymorphism in birds. - Bio.Rev., 79: 815–848. https://doi.org/10.1017/S1464793104006487

Schoenle, L. A., Kernbach, M., Haussmann, M. F., Bonier, F. and Moore, I. 2017. An experimental test of the physiological consequences of avian malaria infection. - J. Anim. Ecol. https://doi.org/10.1111/1365-2656.12753

Schwanz, L. E. 2008. Persistent Effects of Maternal Parasitic Infection on Offspring Fitness: Implications for Adaptive Reproductive Strategies When Parasitized. - Funct. Ecol., 22: 691–698. http://www.jstor.org/stable/20142858

Sheldon, B. C. and Verhulst, S. 1996. Ecological immunology: costly parasite defences and trade-offs in evolutionary ecology. - Trends Ecol. Evol., 11: 317–321. https://doi.org/10.1016/0169-5347(96)10039-2

Thorneycroft, H. B. 1966. Chromosomal polymorphism in the white-throated sparrow, Zonotrichia albicollis (Gmelin). - Science, 154: 1571–1572. https://doi.org/10.1126/science.154.3756.1571

Thorneycroft, H. B. 1975. A cytogenetic study of the white-throated sparrow, Zonotrichia albicollis (Gmelin). - Evolution: 611–621. https://doi.org/10.2307/2407072.

Tuttle, E. M. 2003. Alternative reproductive strategies in the white-throated sparrow: behavioral and genetic evidence. - Behav. Ecol., 14: 425–432. https://doi.org/10.1093/beheco/14.3.425

Tuttle, E. M., Bergland, A. O., Korody, M. L., Brewer, M. S., Newhouse, D. J., Minx, P., Stager, M., Betuel, A., Cheviron, Z. A. and Warren, W. C. 2016. Divergence and functional degradation of a sex chromosome-like supergene. - Curr. Biol., 26: 344–350. https://doi.org/10.1016/j.cub.2015.11.069

Valkiūnas, G. 2004. Avian malaria parasites and other haemosporidia, CRC press.

Valkiūnas, G., Iezhova, T. A. and Shapoval, A. P. 2003. High prevalence of blood parasites in hawfinch Coccothraustes coccothraustes. - J. Nat. Hist., 37: 2647–2652. https://doi.org/10.1080/002229302100001033221

Van Rooyen, J., Lalubin, F., Glaizot, O. and Christe, P. 2013. Avian haemosporidian persistence and co-infection in great tits at the individual level. - Malar. J., 12: 8. https://doi.org/10.1186/1475-2875-12-40

Vedder, O., Komdeur, J., Van Der Velde, M., Schut, E. and Magrath, M. J. L. 2011. Polygyny and extra-pair paternity enhance the opportunity for sexual selection in blue tits. - Behav. Ecol. Sociobiol., 65: 741–752. https://doi.org/10.1007/s00265-010-1078-x

Zehtindjiev, P., Ilieva, M., Westerdahl, H., Hansson, B., Valkiūnas, G. and Bensch, S. 2008. Dynamics of parasitemia of malaria parasites in a naturally and experimentally infected migratory songbird, the great reed warbler Acrocephalus arundinaceus. - Exp. Parasitol., 119: 99–110. https://doi.org/10.1016/j.exppara.2007.12.018

