## Supplementary methods and tables for "Effects of haemosporidian co-infection and parasitemia on reproductive strategies in a polymorphic species"

**SUPPLEMENTARY MATERIALS:**

**Body condition:**

This index was calculated as $Weight\ individual * \left(\frac{Tarsus\ mean\ Population}{Tarsus\ Individual}\right)^{m}$, m being the slope of a linear regression between the weight and the tarsus of the population (Peig and Green 2009, Pigeault et al. 2020). Separate linear regression between weight and tarsus of the population was made for adults and offspring.

**Parentage:**

We based the paternity analysis on seven microsatellites loci: Gf01 (Petren 1998), MME1 (Jeffery et al. 2001), DPμ01 and DPμ03 (Dawson et al. 1997), ZLC02, ZLC07, and ZLH02 (Poesel et al. 2009) with at least four diagnosed microsatellites per individuals. First, we compared the nestling's microsatellites to their parents to determine within- or extra-pair offspring. We assigned them extra-pair if their genotypes did not correspond to the social paternal microsatellites (Grunst et al. 2019). We used CERVUS 3.0.7 (Field Genetics, London, UK, (Kalinowski et al. 2007)) to assign any extra-pair paternity and confirm any within-pair paternity with paternity given at 80% level minimum following the method used in Grunst et al. (2019). Brood parasitism between females is rare in white-throated sparrows (Tuttle 2003). Still, when the social mother had mismatched alleles with its offspring, we manually checked the genotype and attempted to assign the genetic mother. Otherwise, if the social mother had zero mismatches, she was considered the genetic mother. When the social father was not matching the genetic father, we manually compared the genotype of any male with 0-1 mismatch allele and the social father (Formica and Tuttle 2009).

**Parasite determination:**

We used a nested PCR with external primer HaemNFI (5' - CATATATTAAGAGAAITATGGAG – 3') and HaemNR3 (5' - ATAGAAAGATAAGAAATACCATTC – 3') and internal primers HaemF(5'-ATGGTGCTTTCGATATATGCATG-3)/HaemR2(5'-GCATTATCTGGATGTGA TAATGGT-3') specific for *Haemoproteus* and *Plasmodium* (480 bp excluding primer) and a different set of internal primer specific for *Leucocytozoon*: HaemFL (5'-ATGGTGTTTTAGATACTTACATT-3')/HaemR2L (5'-CATTATCTGGATGAGATAATG GIGC-3') (478 bp excluding primer) (Hellgren et al. 2004). We used the same cycling conditions as Hellgren et al. (2004), but numbers of cycles can be added to a total of 40 cycles to ensure the negativity of the possible negative results. We included a negative control (DNA-free water) and positive control in each reaction. The positive sample comes from an individual with a confirmed infection by microscopy. We then ran the PCR product in a 1% agarose gel dyed with ethidium bromide. We repeated the PCR in triplicate to confirm the PCR diagnostic. We then purified all the positive samples using AMPpure XP Beads (ratio 50:50 PCR product diluted in water and beads) to remove potential primer-dimer and salts. Once cleaned, we followed the sequencing PCR protocol (personal communication of H.G.) with 1 µl BigDye Terminator v3.1 ready reaction mix (ThermoFisher Scientific Inc, Waltham, MA USA), 0.5 µl primer, and a combination of 3 µl purified DNA and 3.5 µl of DNA free water for a total solution of 8 µl. The sequencing used the following condition: incubation of 1 min at 94 C°, followed by 25 cycles of 10 sec at 94 C°, 20 sec at 52 C° and 4 min at 72 C°, finishing with 4 C° forever. Once the thermocycling was completed, we cleaned the produce of this reaction with AMPpure XP Beads and sequenced them with SeqStudio sequencer (ThermoFisher Scientific Inc, Waltham, MA USA) with forward and reverse internal primer (HaemF/HaemR2). The PCR products of HeamFL-HeamR2L were sent to MCLAB Inc. (https://www.mclab.com) and sequenced with ABI 3730XL sequencers (ThermoFisher

Scientific Inc, Waltham, MA USA). We trimmed the primers, aligned and corrected the sequences with BioEdit (Hall 1999).

**Parasitemia determination:**

We used the primer R330F (5'- CGTTCTTAACCCAGCTCACG – 3') and R480RL (5'- GCCTGGAGGTWAYGTCC – 3'), amplifying a 182 base pair fragment of Haemosporidian's mitochondrial DNA (Bell et al. 2015a). DNA concentration was measured with NanoDrop$^{TM}$ (ThermoFisher Scientific Inc, Waltham, MA USA), and all samples were diluted to a concentration of 20ng/ul. Reactions were performed using iTaq universal SYBR Green Supermix on QuantStudio (Applied Biosystem, ThermoFisher, Waltham, MA USA). The reaction of a total volume of 10 µl contained 7.5 µl of SYBR Green Supermix, 0.6 µl of each primer (at a 10 µM concentration), 3.3 µl DNA free water, and 2 µl of DNA template (at a concentration of 20 ng/µl) (Bell et al. 2015a). The cycling condition followed the set-up of Bell et al. (2015a). We used DNA-free water as negative control and a synthetic double-stranded DNA product of *Plasmodium relictum* as the positive control (Bell et al. 2015b). We performed all the reactions in triplicate and inspected the melting curve to confirm that the DNA targeted was the one amplified (Bell et al. 2015b). Then, we averaged the parasite concentration between triplicates to reach an average parasite concentration for each sample. We used 4 serial dilutions (1:100 to 1:100000) of the artificial Plasmodium DNA to get a standard curve and quantify the concentration of the parasite in each blood sample. Results were taken into account only when the standard curve slope was not steeper than -3.8, with an efficacy between 82.8% to 97.6% (Asghar et al. 2012) and when the intra- and inter-plate variations were lower than 5% (Bentz et al. 2006).

### SUPPLEMENTARY TABLES

**Table S1:** data description for adults' females, and males with average age and body condition per year, as well as data description of nests for each year with average clutch size, number of fledglings, $G_o/S_f$, and mean nestling body condition and growth rate.

| | | Morph | | |
|---|---|---|---|---|
| **Year** | **Adult Female** | **Tan** | **White** | **adult body condition** |
| 2019 | 17 | 13 | 4 | 24.27±1.21 |
| 2017 | 5 | 0 | 5 | 24.23±1.21 |
| 2016 | 3 | 2 | 1 | 23.37±0.54 |
| 2015 | 2 | 0 | 2 | 23.03±1.52 |
| 2014 | 2 | 1 | 1 | 23.2±0.67 |
| 2013 | 4 | 2 | 2 | 23.67±0.60 |
| 2011 | 2 | 1 | 1 | 23.13±0.4 |
| 2010 | 5 | 3 | 2 | 23.13±1.2 |
| 2007 | 3 | 2 | 1 | 23.01±0.87 |
| 2005 | 1 | 1 | 0 | 21.65±0 |
| 2002 | 1 | 1 | 0 | 20.97±0 |

| | | Morph | | |
|---|---|---|---|---|
| **Year** | **Adult Male** | **Tan** | **White** | **adult body condition** |
| 2019 | 12 | 4 | 8 | 24.35±1.28 |
| 2018 | 1 | 1 | 0 | 26.15±0 |
| 2017 | 5 | 3 | 2 | 24.46±0.69 |
| 2016 | 2 | 1 | 1 | 24.14±0.24 |
| 2015 | 1 | 1 | 0 | 25.31±0 |
| 2014 | 5 | 2 | 3 | 24.83±1 |
| 2013 | 6 | 3 | 3 | 25.26±0.97 |
| 2011 | 1 | 1 | 0 | 22.95±0 |
| 2010 | 5 | 3 | 2 | 24.51±0.51 |
| 2009 | 2 | 1 | 1 | 25.07±0 |
| 2007 | 3 | 1 | 2 | 24.08±0.03 |
| 2005 | 1 | 1 | 0 | NA |
| 2003 | 1 | 0 | 1 | 25.16±0 |

| | | | | Pair Type | | Clutch | | | | | | |
|---|---|---|---|---|---|---|---|---|---|---|---|---|
| **Year** | **Nest** | **Father** | **Mother** | **T×W** | **W×T** | **1** | **2** | **Clutch size** | **nb fledglings** | **$G_o/S_f$** | **Nestling body condition** | **Nestling Growth rate** |
| 2019 | 26 | 16 | 25 | 7 | 19 | 19 | 6 | 2.96±0.91 | 2.04±1.79 | 0.58±0.43 | 13.45±0.77 | 2.31±0.2 |
| 2018 | 2 | 2 | 0 | 1 | 1 | 2 | 0 | 3.5±0.71 | 1.5±1.12 | 1±0 | 14.66±1 | 2.38±0.06 |
| 2017 | 11 | 10 | 8 | 8 | 2 | 8 | 3 | 3.55±1.21 | 2.27±2.05 | 0.82±0.40 | 12.77±0.89 | 2.18±0.23 |
| 2016 | 7 | 3 | 5 | 2 | 4 | 5 | 1 | 2.83±1.33 | 2.17±2.04 | 0.75±0.29 | 12.98±0.97 | 2.79±0.28 |
| 2015 | 4 | 2 | 2 | 4 | 0 | 2 | 2 | 3.75±1.26 | 3.5±1.29 | 0.92±0.14 | 12.85±0.8 | 2.2±0.33 |
| 2014 | 8 | 7 | 2 | 3 | 5 | 7 | 1 | 3.75±0.89 | 2.75±2.05 | 0.875±0.35 | 12.87±0.84 | 2.33±0.3 |
| 2013 | 11 | 9 | 7 | 6 | 5 | 8 | 3 | 2.72±1.19 | 2.18±1.6 | 0.85±0.35 | 12.6±0.7 | 2.39±0.27 |
| 2011 | 2 | 1 | 2 | 1 | 1 | 2 | 0 | 3.5±0.7 | 3.5±0.7 | 0.83±0.24 | 14.74±0.08 | 2.60±0.04 |
| 2010 | 8 | 6 | 7 | 3 | 5 | 6 | 1 | 2.86±1.35 | 2.43±1.13 | 0.43±0.4 | 13.12±1.2 | 2.11±0.45 |
| 2009 | 3 | 3 | 0 | 2 | 1 | 2 | 1 | 3.33±1.15 | 3.33±1.15 | 1±0 | 13.66±0.52 | 2.59±0.24 |
| 2007 | 5 | 3 | 4 | 3 | 2 | 4 | 1 | 4±0 | 4±0 | 1±0 | 13.8±0.92 | 2.27±0.25 |
| 2005 | 2 | 1 | 1 | 1 | 1 | 2 | 0 | 4±0 | 4±0 | 1±0 | 12.5±0.71 | 1.65±0.48 |

| | | | | | | | | | | | | |
|---|---|---|---|---|---|---|---|---|---|---|---|---|
| 2003 | 1 | 1 | 0 | 0 | 1 | 1 | 0 | 4±0 | 4±0 | 1±0 | 13.66±0.48 | 2.28±0.14 |
| 2002 | 1 | 0 | 1 | 0 | 1 | 1 | 0 | 3±0 | 3±0 | 1±0 | 15.57±0.94 | 1.48±0.62 |

**Table S1 cont':**

Table S2a: Models on the effect sex morph and population density on adult's body condition with coinfection. $\omega$ is the Akaike weight.

| Models | Morph (white) | population density | Coinfection | Sex (sex) | White morph×male | df | AICc | $\Delta$AICc | $\omega$ |
|---|---|---|---|---|---|---|---|---|---|
| 15 | | × | × | × | | 6 | 227.32 | 0.00 | 0.301 |
| 11 | | × | | × | | 5 | 228.02 | 0.70 | 0.212 |
| 13 | | | × | × | | 5 | 229.02 | 1.70 | 0.129 |
| 9 | | | | × | | 4 | 229.41 | 2.09 | 0.106 |
| 16 | × | × | × | × | | 7 | 230.51 | 3.19 | 0.061 |
| 12 | × | × | | × | | 6 | 231.13 | 3.81 | 0.045 |
| 32 | × | × | × | × | × | 8 | 231.89 | 4.57 | 0.031 |
| 14 | × | | × | × | | 6 | 232.19 | 4.87 | 0.026 |
| 28 | × | × | | × | × | 7 | 232.43 | 5.11 | 0.023 |
| 10 | × | | | × | | 5 | 232.49 | 5.17 | 0.023 |
| 30 | × | | × | × | × | 7 | 232.85 | 5.53 | 0.019 |
| 26 | × | | | × | × | 6 | 233.29 | 5.97 | 0.015 |
| 3 | | × | | | | 4 | 236.60 | 9.28 | 0.003 |
| 7 | | × | × | | | 5 | 237.67 | 10.35 | 0.002 |
| 1 | | | | | | 3 | 237.71 | 10.39 | 0.002 |
| 5 | | | × | | | 4 | 238.89 | 11.57 | 0.001 |
| 4 | × | × | | | | 5 | 239.48 | 12.16 | 0.001 |

| | | | | | | | | | |
|---|---|---|---|---|---|---|---|---|---|
| 2 | × | | | | | 4 | 240.50 | 13.18 | 0.000 |
| 8 | × | × | × | | | 6 | 240.65 | 13.33 | 0.000 |
| 6 | × | | × | | | 5 | 241.77 | 14.45 | 0.000 |

Table S2b: Models on the effect sex morph and population density on adult's body condition with parasitemia. *ω* is the Akaike weight.

| **Models** | **Morph (Sex)** | **Parasitemia** | **population density** | **Sex (male)** | **White morph×male** | **df** | **AICc** | ***Δ*AICc** | ***ω*** |
|---|---|---|---|---|---|---|---|---|---|
| 15 | | × | × | × | | 6 | 222.62 | 0.00 | 0.483 |
| 11 | | × | | × | | 5 | 224.21 | 1.59 | 0.218 |
| 16 | × | × | × | × | | 7 | 225.77 | 3.14 | 0.100 |
| 32 | × | × | × | × | × | 8 | 227.22 | 4.6 | 0.048 |
| 12 | × | × | | × | | 6 | 227.34 | 4.72 | 0.046 |
| 13 | | | × | × | | 5 | 228.02 | 5.39 | 0.033 |
| 28 | × | × | | × | × | 7 | 228.23 | 5.61 | 0.029 |
| 9 | | | | × | | 4 | 229.41 | 6.78 | 0.016 |
| 14 | × | | × | × | | 6 | 231.13 | 8.51 | 0.007 |
| 7 | | × | × | | | 5 | 231.61 | 8.99 | 0.005 |
| 30 | × | | × | × | × | 7 | 232.43 | 9.81 | 0.004 |
| 10 | × | | | × | | 5 | 232.49 | 9.86 | 0.003 |
| 3 | | × | | | | 4 | 232.73 | 10.1 | 0.003 |
| 26 | × | | | × | × | 6 | 233.29 | 10.67 | 0.002 |
| 8 | × | × | × | | | 6 | 234.56 | 11.94 | 0.001 |
| 4 | × | × | | | | 5 | 235.57 | 12.95 | 0.001 |
| 5 | | | × | | | 4 | 236.60 | 13.97 | 0.000 |

| | | | | | | | | | |
|---|---|---|---|---|---|---|---|---|---|
| 1 | | | | | | 3 | 237.71 | 15.09 | 0.000 |
| 6 | × | | × | | | 5 | 239.48 | 16.86 | 0.000 |
| 2 | × | | | | | 4 | 240.50 | 17.88 | 0.000 |

Table S3a: Models on the effect of clutch order and mothers' body condition, morph, co-infection status, and parasitemia on fledgling numbers. $\omega$ is the Akaike weight.

| **Models** | **Clutch (second)** | **Mother body condition** | **Mother co-infection** | **Mother morph (white)** | **Mother parasitemia** | **df** | **AICc** | ***Δ*AICc** | ***ω*** |
|---|---|---|---|---|---|---|---|---|---|
| 11 | | × | | × | | 4 | 198.58 | 0.00 | 0.101 |
| 3 | | × | | | | 3 | 198.60 | 0.02 | 0.100 |
| 4 | × | × | | | | 4 | 198.74 | 0.15 | 0.094 |
| 12 | × | × | | × | | 5 | 198.86 | 0.27 | 0.088 |
| 2 | × | | | | | 3 | 199.42 | 0.83 | 0.067 |
| 10 | × | | | × | | 4 | 199.95 | 1.37 | 0.051 |
| 1 | | | | | | 2 | 200.28 | 1.69 | 0.043 |
| 9 | | | | × | | 3 | 200.72 | 2.14 | 0.035 |
| 7 | | × | × | | | 4 | 200.80 | 2.21 | 0.034 |
| 8 | × | × | × | | | 5 | 200.87 | 2.29 | 0.032 |
| 15 | | × | × | × | | 5 | 200.97 | 2.38 | 0.031 |
| 19 | | × | | | × | 4 | 200.97 | 2.39 | 0.031 |
| 27 | | × | | × | × | 5 | 201.03 | 2.45 | 0.030 |
| 20 | × | × | | | × | 5 | 201.18 | 2.59 | 0.028 |

| | | | | | | | | | |
|---|---|---|---|---|---|---|---|---|---|
| 16 | × | × | × | × | | 6 | 201.24 | 2.66 | 0.027 |
| 28 | × | × | | × | × | 6 | 201.45 | 2.86 | 0.024 |
| 6 | × | | × | | | 4 | 201.68 | 3.09 | 0.022 |
| 18 | × | | | | × | 4 | 201.75 | 3.16 | 0.021 |
| 26 | × | | | × | × | 5 | 202.24 | 3.65 | 0.016 |
| 17 | | | | | × | 3 | 202.32 | 3.73 | 0.016 |
| 14 | × | | × | × | | 5 | 202.40 | 3.82 | 0.015 |
| 5 | | | × | | | 3 | 202.54 | 3.95 | 0.014 |
| 25 | | | | × | × | 4 | 202.57 | 3.99 | 0.014 |
| 13 | | | × | × | | 4 | 203.08 | 4.50 | 0.011 |
| 23 | | × | × | | × | 5 | 203.25 | 4.67 | 0.010 |
| 24 | × | × | × | | × | 6 | 203.46 | 4.88 | 0.009 |
| 31 | | × | × | × | × | 6 | 203.47 | 4.88 | 0.009 |
| 32 | × | × | × | × | × | 7 | 203.92 | 5.34 | 0.007 |
| 22 | × | | × | | × | 5 | 204.06 | 5.47 | 0.007 |
| 21 | | | × | | × | 4 | 204.62 | 6.04 | 0.005 |
| 30 | × | | × | × | × | 6 | 204.71 | 6.12 | 0.005 |
| 29 | | | × | × | × | 5 | 205.00 | 6.42 | 0.004 |

Table S3b: Models on the effect of clutch order and fathers' body condition, morph, co-infection status and parasitemia on fledgling numbers. *ω* is the Akaike weight.

| Models | Clutch (second) | Father body condition | Father co-infection | Father morph (white) | Father parasitemia | df | AICc | *Δ*AICc | *ω* |
|---|---|---|---|---|---|---|---|---|---|
| 18 | × | | | | × | 4 | 156.0 | 0.0 | 0.101 |
| 2 | × | | | | | 3 | 156.2 | 0.1 | 0.095 |
| 24 | × | × | × | | × | 6 | 156.7 | 0.7 | 0.072 |
| 10 | × | | | × | | 4 | 156.9 | 0.9 | 0.066 |
| 22 | × | | × | | × | 5 | 157.1 | 1.0 | 0.060 |
| 32 | × | × | × | × | × | 7 | 157.4 | 1.4 | 0.051 |
| 20 | × | × | | | × | 5 | 157.4 | 1.4 | 0.050 |
| 26 | × | | | × | × | 5 | 157.5 | 1.4 | 0.050 |
| 4 | × | × | | | | 4 | 157.6 | 1.5 | 0.047 |
| 12 | × | × | | × | | 5 | 158.0 | 2.0 | 0.038 |
| 6 | × | | × | | | 4 | 158.2 | 2.1 | 0.035 |
| 17 | | | | | × | 3 | 158.3 | 2.3 | 0.032 |
| 28 | × | × | | × | × | 6 | 158.6 | 2.6 | 0.028 |
| 30 | × | | × | × | × | 6 | 158.7 | 2.6 | 0.027 |
| 16 | × | × | × | × | | 6 | 158.8 | 2.7 | 0.026 |
| 8 | × | × | × | | | 5 | 158.8 | 2.7 | 0.026 |

| | | | | | | | | | |
|---|---|---|---|---|---|---|---|---|---|
| 14 | × | | × | × | | 5 | 159.0 | 2.9 | 0.023 |
| 25 | | | | × | × | 4 | 159.3 | 3.3 | 0.020 |
| 21 | | | × | | × | 4 | 159.4 | 3.3 | 0.019 |
| 1 | | | | | | 2 | 159.4 | 3.4 | 0.019 |
| 9 | | | | × | | 3 | 159.5 | 3.5 | 0.018 |
| 19 | | × | | | × | 4 | 160.2 | 4.1 | 0.013 |
| 31 | | × | × | × | × | 6 | 160.2 | 4.1 | 0.013 |
| 23 | | × | × | | × | 5 | 160.3 | 4.2 | 0.012 |
| 29 | | | × | × | × | 5 | 160.4 | 4.3 | 0.012 |
| 27 | | × | | × | × | 5 | 160.9 | 4.8 | 0.009 |
| 11 | | × | | × | | 4 | 161.0 | 5.0 | 0.009 |
| 3 | | × | | | | 3 | 161.3 | 5.2 | 0.007 |
| 5 | | | × | | | 3 | 161.5 | 5.4 | 0.007 |
| 13 | | | × | × | | 4 | 161.5 | 5.4 | 0.007 |
| 15 | | × | × | × | | 5 | 162.1 | 6.0 | 0.005 |
| 7 | | × | × | | | 4 | 163.0 | 7.0 | 0.003 |

Table S4a: Models on the effect of clutch order and mothers' body condition, morph, co-infection status and parasitemia on chances to acquire extra-pair paternity. *ω* is the Akaike weight.

| **Models** | **Clutch (second)** | **Mother body condition** | **Mother co-infection** | **Mother morph (white)** | **Mother parasitemia** | **df** | **AICc** | ***Δ*AICc** | ***ω*** |
|---|---|---|---|---|---|---|---|---|---|
| 30 | × | | × | × | × | 6 | 60.43 | 0.00 | 0.220 |
| 14 | × | | × | × | | 5 | 61.47 | 1.04 | 0.131 |
| 29 | | | × | × | × | 5 | 62.10 | 1.67 | 0.096 |
| 22 | × | | × | | × | 5 | 62.67 | 2.24 | 0.072 |
| 32 | × | × | × | × | × | 7 | 62.94 | 2.50 | 0.063 |
| 21 | | | × | | × | 4 | 63.49 | 3.06 | 0.048 |
| 13 | | | × | × | | 4 | 63.72 | 3.29 | 0.043 |
| 16 | × | × | × | × | | 6 | 64.02 | 3.59 | 0.037 |
| 10 | × | | | × | | 4 | 64.14 | 3.71 | 0.035 |
| 25 | | | | × | × | 4 | 64.50 | 4.07 | 0.029 |
| 31 | | × | × | × | × | 6 | 64.68 | 4.25 | 0.026 |
| 9 | | | | × | | 3 | 64.79 | 4.36 | 0.025 |
| 26 | × | | | × | × | 5 | 64.82 | 4.39 | 0.025 |

| | | | | | | | | | |
|---|---|---|---|---|---|---|---|---|---|
| 24 | × | × | × | | × | 6 | 65.21 | 4.78 | 0.020 |
| 23 | | × | × | | × | 5 | 65.96 | 5.52 | 0.014 |
| 17 | | | | | × | 3 | 66.05 | 5.62 | 0.013 |
| 15 | | × | × | × | | 5 | 66.18 | 5.75 | 0.012 |
| 12 | × | × | | × | | 5 | 66.41 | 5.97 | 0.011 |
| 6 | × | | × | | | 4 | 66.48 | 6.05 | 0.011 |
| 27 | | × | | × | × | 5 | 66.82 | 6.39 | 0.009 |
| 18 | × | | | | × | 4 | 66.83 | 6.39 | 0.009 |
| 28 | × | × | | × | × | 6 | 67.04 | 6.61 | 0.008 |
| 11 | | × | | × | | 4 | 67.14 | 6.71 | 0.008 |
| 2 | × | | | | | 3 | 67.27 | 6.84 | 0.007 |
| 1 | | | | | | 2 | 67.52 | 7.09 | 0.006 |
| 5 | | | × | | | 3 | 67.81 | 7.38 | 0.006 |
| 19 | | × | | | × | 4 | 68.23 | 7.80 | 0.004 |
| 8 | × | × | × | | | 5 | 68.93 | 8.49 | 0.003 |
| 20 | × | × | | | × | 5 | 68.97 | 8.54 | 0.003 |
| 4 | × | × | | | | 4 | 69.61 | 9.18 | 0.002 |
| 3 | | × | | | | 3 | 69.79 | 9.36 | 0.002 |
| 7 | | × | × | | | 4 | 70.02 | 9.58 | 0.002 |

Table S4b: Models on the effect of clutch order and fathers' body condition, morph, co-infection status and parasitemia on chances to acquire extra-pair paternity. $\omega$ is the Akaike weight.

| Models | Clutch (second) | Father body condition | Father co-infection | Father morph (white) | Father Parasitemia | df | AICc | ΔAICc | $\omega$ |
|---|---|---|---|---|---|---|---|---|---|
| 13 | | | × | × | | 4 | 41.90 | 0.00 | 0.206 |
| 9 | | | | × | | 3 | 42.88 | 0.98 | 0.126 |
| 14 | × | | × | × | | 5 | 43.44 | 1.54 | 0.095 |
| 10 | × | | | × | | 4 | 44.24 | 2.34 | 0.064 |
| 11 | | × | | × | | 4 | 44.48 | 2.58 | 0.057 |
| 15 | | × | × | × | | 5 | 44.49 | 2.59 | 0.056 |
| 29 | | | × | × | × | 5 | 44.51 | 2.61 | 0.056 |
| 5 | | | × | | | 3 | 44.81 | 2.91 | 0.048 |
| 25 | | | | × | × | 4 | 45.19 | 3.30 | 0.040 |
| 12 | × | × | | × | | 5 | 45.97 | 4.07 | 0.027 |
| 30 | × | | × | × | × | 6 | 46.20 | 4.30 | 0.024 |
| 16 | × | × | × | × | | 6 | 46.24 | 4.34 | 0.024 |
| 1 | | | | | | 2 | 46.47 | 4.57 | 0.021 |
| 26 | × | | | × | × | 5 | 46.60 | 4.70 | 0.020 |

| | | | | | | | | | |
|---|---|---|---|---|---|---|---|---|---|
| 27 | | × | | × | × | 5 | 46.99 | 5.09 | 0.016 |
| 6 | × | | × | | | 4 | 47.00 | 5.10 | 0.016 |
| 21 | | | × | | × | 4 | 47.08 | 5.18 | 0.015 |
| 7 | | × | × | | | 4 | 47.11 | 5.21 | 0.015 |
| 31 | | × | × | × | × | 6 | 47.27 | 5.37 | 0.014 |
| 17 | | | | | × | 3 | 48.00 | 6.10 | 0.010 |
| 28 | × | × | | × | × | 6 | 48.56 | 6.66 | 0.007 |
| 2 | × | | | | | 3 | 48.65 | 6.75 | 0.007 |
| 3 | | × | | | | 3 | 48.70 | 6.80 | 0.007 |
| 32 | × | × | × | × | × | 7 | 49.18 | 7.28 | 0.005 |
| 8 | × | × | × | | | 5 | 49.27 | 7.37 | 0.005 |
| 22 | × | | × | | × | 5 | 49.34 | 7.44 | 0.005 |
| 23 | | × | × | | × | 5 | 49.58 | 7.68 | 0.004 |
| 18 | × | | | | × | 4 | 50.13 | 8.23 | 0.003 |
| 19 | | × | | | × | 4 | 50.32 | 8.42 | 0.003 |
| 4 | × | × | | | | 4 | 51.06 | 9.16 | 0.002 |
| 24 | × | × | × | | × | 6 | 51.83 | 9.94 | 0.001 |
| 20 | × | × | | | × | 5 | 52.67 | 10.77 | 0.001 |

Table S5a: Models on the effect of clutch order and mothers' body condition, morph, co-infection status and parasitemia on the ratio of genetic offspring belonging to the social father. *ω* is the Akaike weight.

| **Models** | **Clutch (second)** | **Mother body condition** | **Mother co-infection** | **Mother morph (white)** | **Mother parasitemia** | **df** | **AICc** | ***Δ*AICc** | ***ω*** |
|---|---|---|---|---|---|---|---|---|---|
| 10 | × | | | × | | 4 | 133.70 | 0.00 | 0.269 |
| 12 | × | × | | × | | 5 | 134.44 | 0.74 | 0.186 |
| 14 | × | | × | × | | 5 | 135.49 | 1.79 | 0.110 |
| 26 | × | | | × | × | 5 | 136.17 | 2.47 | 0.078 |
| 9 | | | | × | | 3 | 136.75 | 3.05 | 0.059 |
| 16 | × | × | × | × | | 6 | 136.79 | 3.10 | 0.057 |
| 28 | × | × | | × | × | 6 | 137.09 | 3.39 | 0.049 |
| 30 | × | | × | × | × | 6 | 138.14 | 4.44 | 0.029 |
| 11 | | × | | × | | 4 | 138.24 | 4.55 | 0.028 |
| 13 | | | × | × | | 4 | 138.83 | 5.14 | 0.021 |
| 25 | | | | × | × | 4 | 138.94 | 5.25 | 0.020 |
| 2 | × | | | | | 3 | 139.15 | 5.45 | 0.018 |
| 32 | × | × | × | × | × | 7 | 139.57 | 5.87 | 0.014 |

| | | | | | | | | | |
|---|---|---|---|---|---|---|---|---|---|
| 27 | | × | | × | × | 5 | 140.34 | 6.64 | 0.010 |
| 15 | | × | × | × | | 5 | 140.68 | 6.98 | 0.008 |
| 29 | | | × | × | × | 5 | 140.88 | 7.19 | 0.007 |
| 18 | × | | | | × | 4 | 141.08 | 7.38 | 0.007 |
| 4 | × | × | | | | 4 | 141.21 | 7.52 | 0.006 |
| 6 | × | | × | | | 4 | 141.43 | 7.73 | 0.006 |
| 31 | | × | × | × | × | 6 | 142.72 | 9.02 | 0.003 |
| 20 | × | × | | | × | 5 | 142.98 | 9.29 | 0.003 |
| 22 | × | | × | | × | 5 | 143.10 | 9.40 | 0.002 |
| 1 | | | | | | 2 | 143.39 | 9.70 | 0.002 |
| 17 | | | | | × | 3 | 143.45 | 9.75 | 0.002 |
| 8 | × | × | × | | | 5 | 143.70 | 10.00 | 0.002 |
| 24 | × | × | × | | × | 6 | 145.32 | 11.62 | 0.001 |
| 19 | | × | | | × | 4 | 145.48 | 11.78 | 0.001 |
| 21 | | | × | | × | 4 | 145.48 | 11.78 | 0.001 |
| 3 | | × | | | | 3 | 145.67 | 11.98 | 0.001 |
| 5 | | | × | | | 3 | 145.68 | 11.98 | 0.001 |
| 23 | | × | × | | × | 5 | 147.76 | 14.06 | 0.000 |
| 7 | | × | × | | | 4 | 148.07 | 14.37 | 0.000 |

Table S5b: Models on the effect of clutch order and fathers' body condition, morph, co-infection status and parasitemia on the ratio of genetic offspring belonging to the social father. $\omega$ is the Akaike weight.

| Models | Clutch (second) | Father body condition | Father co-infection | Father morph (white) | Father parasitemia | df | AICc | $\Delta$AICc | $\omega$ |
|---|---|---|---|---|---|---|---|---|---|
| 29 | | | × | × | × | 5 | 83.01 | 0.00 | 0.296 |
| 25 | | | | × | × | 4 | 83.27 | 0.26 | 0.260 |
| 30 | × | | × | × | × | 6 | 85.27 | 2.26 | 0.096 |
| 26 | × | | | × | × | 5 | 85.54 | 2.53 | 0.083 |
| 27 | | × | | × | × | 5 | 85.70 | 2.69 | 0.077 |
| 31 | | × | × | × | × | 6 | 85.76 | 2.75 | 0.075 |
| 28 | × | × | | × | × | 6 | 88.10 | 5.09 | 0.023 |
| 32 | × | × | × | × | × | 7 | 88.21 | 5.19 | 0.022 |
| 9 | | | | × | | 3 | 88.24 | 5.23 | 0.022 |
| 10 | × | | | × | | 4 | 88.56 | 5.55 | 0.018 |
| 13 | | | × | × | | 4 | 90.54 | 7.53 | 0.007 |
| 11 | | × | | × | | 4 | 90.65 | 7.64 | 0.006 |
| 14 | × | | × | × | | 5 | 90.76 | 7.75 | 0.006 |
| 12 | × | × | | × | | 5 | 91.06 | 8.05 | 0.005 |

| | | | | | | | | | |
|---|---|---|---|---|---|---|---|---|---|
| 15 | | × | × | × | | 5 | 93.18 | 10.17 | 0.002 |
| 16 | × | × | × | × | | 6 | 93.56 | 10.55 | 0.002 |
| 23 | | × | × | | × | 5 | 108.09 | 25.08 | 0.000 |
| 24 | × | × | × | | × | 6 | 110.90 | 27.89 | 0.000 |
| 7 | | × | × | | | 4 | 111.12 | 28.11 | 0.000 |
| 21 | | | × | | × | 4 | 111.45 | 28.44 | 0.000 |
| 5 | | | × | | | 3 | 112.13 | 29.12 | 0.000 |
| 1 | | | | | | 2 | 112.98 | 29.97 | 0.000 |
| 8 | × | × | × | | | 5 | 113.51 | 30.50 | 0.000 |
| 22 | × | | × | | × | 5 | 114.06 | 31.05 | 0.000 |
| 6 | × | | × | | | 4 | 114.34 | 31.33 | 0.000 |
| 17 | | | | | × | 3 | 114.67 | 31.66 | 0.000 |
| 3 | | × | | | | 3 | 114.83 | 31.82 | 0.000 |
| 2 | × | | | | | 3 | 115.22 | 32.21 | 0.000 |
| 19 | | × | | | × | 4 | 116.62 | 33.61 | 0.000 |
| 18 | × | | | | × | 4 | 117.15 | 34.14 | 0.000 |
| 4 | × | × | | | | 4 | 117.21 | 34.20 | 0.000 |
| 20 | × | × | | | × | 5 | 119.25 | 36.24 | 0.000 |

Table S6a: Models on the effect of clutch order and mothers' body condition, morph, co-infection status and parasitemia on nestling growth rate. $\omega$ is the Akaike weight.

| Models | Clutch (second) | Mother Body Condition | Mother co-infection | Mother morph (white) | Mother Parasitemia | df | AICc | $\Delta$AICc | $\omega$ |
|---|---|---|---|---|---|---|---|---|---|
| 17 | | | | | × | **4** | 48.49055 | 0 | 0.245 |
| 18 | × | | | | × | **5** | 48.79093 | 0.300374 | 0.211 |
| 22 | × | | × | | × | **6** | 49.50114 | 1.010584 | 0.148 |
| 21 | | | × | | × | **5** | 50.26042 | 1.769863 | 0.101 |
| 1 | | | | | | **3** | 51.27384 | 2.783291 | 0.061 |
| 2 | × | | | | | **4** | 51.77914 | 3.288583 | 0.047 |
| 6 | × | | × | | | **5** | 51.97108 | 3.480531 | 0.043 |
| 5 | | | × | | | **4** | 53.04897 | 4.558421 | 0.025 |
| 25 | | | | × | × | **5** | 53.18345 | 4.692895 | 0.023 |
| 26 | × | | | × | × | **6** | 53.60593 | 5.11538 | 0.019 |
| 30 | × | | × | × | × | **7** | 54.28401 | 5.793459 | 0.014 |
| 29 | | | × | × | × | **6** | 54.95434 | 6.463784 | 0.010 |
| 19 | | × | | | × | **5** | 55.00765 | 6.517098 | 0.009 |
| 20 | × | × | | | × | **6** | 55.03424 | 6.543685 | 0.009 |
| 9 | | | | × | | **4** | 55.9438 | 7.453251 | 0.006 |
| 24 | × | × | × | | × | **7** | 56.37089 | 7.880337 | 0.005 |

| | | | | | | | | | |
|---|---|---|---|---|---|---|---|---|---|
| 10 | × | | | × | | **5** | 56.64177 | 8.151222 | 0.004 |
| 14 | × | | × | × | | **6** | 56.66929 | 8.178741 | 0.004 |
| 23 | | × | × | | × | **6** | 57.20696 | 8.716405 | 0.003 |
| 13 | | | × | × | | **5** | 57.59131 | 9.100754 | 0.003 |
| 4 | × | × | | | | **5** | 57.70318 | 9.212627 | 0.002 |
| 3 | | × | | | | **4** | 57.72355 | 9.232995 | 0.002 |
| 8 | × | × | × | | | **6** | 58.78548 | 10.292928 | 0.001 |
| 27 | | × | | × | × | **6** | 59.76471 | 11.27416 | 0.001 |
| 28 | × | × | | × | × | **7** | 59.9006 | 11.410048 | 0.001 |
| 7 | | × | × | | | **5** | 60.01165 | 11.521096 | 0.001 |
| 32 | × | × | × | × | × | **8** | 61.23122 | 12.740666 | 0.000 |
| 31 | | × | × | × | × | **7** | 61.99029 | 13.499738 | 0.000 |
| 11 | | × | | × | | **5** | 62.40493 | 13.914382 | 0.000 |
| 12 | × | × | | × | | **6** | 62.55252 | 14.061967 | 0.000 |
| 16 | × | × | × | × | | **7** | 63.51437 | 15.023823 | 0.000 |
| 15 | | × | × | × | | **6** | 64.61863 | 16.128081 | 0.000 |

Table S6b: Models on the effect of clutch order and father' body condition, morph, co-infection status and parasitemia on nestling growth rate. $\omega$ is the Akaike weight.

| Models | Clutch (second) | Father body condition | Father co-infection | Father morph (white) | Father Parasitemia | df | AICc | $\Delta$AICc | $\omega$ |
|---|---|---|---|---|---|---|---|---|---|
| 17 | | | | | × | 4 | 19.84 | 0.00 | 0.393 |
| 18 | × | | | | × | 5 | 21.09 | 1.24 | 0.211 |
| 25 | | | | × | × | 5 | 23.12 | 3.27 | 0.077 |
| 21 | | | × | | × | 5 | 23.68 | 3.84 | 0.058 |
| 1 | | | | | | 3 | 23.71 | 3.86 | 0.057 |
| 2 | × | | | | | 4 | 24.03 | 4.19 | 0.048 |
| 26 | × | | | × | × | 6 | 24.80 | 4.96 | 0.033 |
| 22 | × | | × | | × | 6 | 24.95 | 5.11 | 0.031 |
| 19 | | × | | | × | 5 | 26.16 | 6.32 | 0.017 |
| 9 | | | | × | | 4 | 26.20 | 6.35 | 0.016 |
| 29 | | | × | × | × | 6 | 27.12 | 7.27 | 0.010 |
| 10 | × | | | × | | 5 | 27.12 | 7.27 | 0.010 |
| 20 | × | × | | | × | 6 | 27.98 | 8.13 | 0.007 |
| 5 | | | × | | | 4 | 28.20 | 8.36 | 0.006 |
| 6 | × | | × | | | 5 | 28.47 | 8.63 | 0.005 |
| 30 | × | | × | × | × | 7 | 28.85 | 9.01 | 0.004 |

| | | | | | | | | | |
|---|---|---|---|---|---|---|---|---|---|
| 3 | | × | | | | 4 | 29.64 | 9.79 | 0.003 |
| 27 | | × | | × | × | 6 | 30.00 | 10.15 | 0.002 |
| 23 | | × | × | | × | 6 | 30.52 | 10.67 | 0.002 |
| 4 | × | × | | | | 5 | 30.66 | 10.81 | 0.002 |
| 13 | | | × | × | | 5 | 30.75 | 10.90 | 0.002 |
| 14 | × | | × | × | | 6 | 31.62 | 11.78 | 0.001 |
| 28 | × | × | | × | × | 7 | 32.05 | 12.21 | 0.001 |
| 24 | × | × | × | | × | 7 | 32.12 | 12.28 | 0.001 |
| 11 | | × | | × | | 5 | 32.84 | 12.99 | 0.001 |
| 12 | × | × | | × | | 6 | 34.17 | 14.32 | 0.000 |
| 31 | | × | × | × | × | 7 | 34.18 | 14.33 | 0.000 |
| 7 | | × | × | | | 5 | 34.52 | 14.67 | 0.000 |
| 8 | × | × | × | | | 6 | 35.40 | 15.56 | 0.000 |
| 32 | × | × | × | × | × | 8 | 36.01 | 16.17 | 0.000 |
| 15 | | × | × | × | | 6 | 37.61 | 17.77 | 0.000 |
| 16 | × | × | × | × | | 7 | 38.71 | 18.87 | 0.000 |

Table S7a: Models on the effect of clutch order and mothers' body condition, morph, co-infection status and parasitemia on nestling body condition. *ω* is the Akaike weight.

| **Models** | **Clutch (second)** | **Mother Body Condition** | **Mother co-infection** | **Mother morph (white)** | **Mother Parasitemia** | **df** | **AICc** | ***ΔAICc*** | ***ω*** |
|---|---|---|---|---|---|---|---|---|---|
| 22 | × | | × | | × | 6 | 140.39 | 0.00 | 0.469 |
| 21 | | | × | | × | 5 | 142.68 | 2.29 | 0.149 |
| 24 | × | × | × | | × | 7 | 143.20 | 2.82 | 0.115 |
| 30 | × | | × | × | × | 7 | 143.29 | 2.90 | 0.110 |
| 29 | | | × | × | × | 6 | 145.54 | 5.15 | 0.036 |
| 23 | | × | × | | × | 6 | 146.01 | 5.62 | 0.028 |
| 32 | × | × | × | × | × | 8 | 146.04 | 5.66 | 0.028 |
| 18 | × | | | | × | 5 | 147.63 | 7.24 | 0.013 |
| 6 | × | | × | | | 5 | 147.74 | 7.35 | 0.012 |
| 17 | | | | | × | 4 | 148.52 | 8.13 | 0.008 |
| 31 | | × | × | × | × | 7 | 148.90 | 8.51 | 0.007 |
| 20 | × | × | | | × | 6 | 149.54 | 9.15 | 0.005 |
| 14 | × | | × | × | | 6 | 149.97 | 9.59 | 0.004 |
| 26 | × | | | × | × | 6 | 150.26 | 9.87 | 0.003 |
| 19 | | × | | | × | 5 | 150.76 | 10.37 | 0.003 |

| | | | | | | | | | |
|---|---|---|---|---|---|---|---|---|---|
| 25 | | | | × | × | 5 | 151.09 | 10.70 | 0.002 |
| 8 | × | × | × | | | 6 | 151.93 | 11.54 | 0.001 |
| 5 | | | × | | | 4 | 152.00 | 11.61 | 0.001 |
| 28 | × | × | | × | × | 7 | 152.09 | 11.71 | 0.001 |
| 2 | × | | | | | 4 | 152.39 | 12.00 | 0.001 |
| 27 | | × | | × | × | 6 | 153.34 | 12.95 | 0.001 |
| 16 | × | × | × | × | | 7 | 153.93 | 13.55 | 0.001 |
| 13 | | | × | × | | 5 | 154.07 | 13.68 | 0.001 |
| 1 | | | | | | 3 | 154.13 | 13.75 | 0.000 |
| 10 | × | | | × | | 5 | 154.90 | 14.52 | 0.000 |
| 4 | × | × | | | | 5 | 155.23 | 14.84 | 0.000 |
| 9 | | | | × | | 4 | 156.42 | 16.03 | 0.000 |
| 7 | | × | × | | | 5 | 156.56 | 16.17 | 0.000 |
| 12 | × | × | | × | | 6 | 157.38 | 16.99 | 0.000 |
| 3 | | × | | | | 4 | 157.78 | 17.40 | 0.000 |
| 15 | | × | × | × | | 6 | 158.57 | 18.18 | 0.000 |
| 11 | | × | | × | | 5 | 159.85 | 19.46 | 0.000 |

Table S7b: Models on the effect of clutch order and fathers' body condition, morph, co-infection status and parasitemia on nestling body condition. *ω* is the Akaike weight.

| **Models** | **Clutch (second)** | **Father body condition** | **Father co-infection** | **Father morph (white)** | **Father Parasitemia** | **df** | **AICc** | ***Δ*AICc** | ***ω*** |
|---|---|---|---|---|---|---|---|---|---|
| 25 | | | | × | × | 5 | 78.96 | 0.00 | 0.442 |
| 27 | | × | | × | × | 6 | 81.78 | 2.82 | 0.108 |
| 29 | | | × | × | × | 6 | 81.82 | 2.86 | 0.106 |
| 26 | × | | | × | × | 6 | 81.92 | 2.96 | 0.101 |
| 9 | | | | × | | 4 | 82.82 | 3.86 | 0.064 |
| 30 | × | | × | × | × | 7 | 85.04 | 6.08 | 0.021 |
| 28 | × | × | | × | × | 7 | 85.07 | 6.11 | 0.021 |
| 11 | | × | | × | | 5 | 85.27 | 6.31 | 0.019 |
| 13 | | | × | × | | 5 | 85.31 | 6.35 | 0.018 |
| 31 | | × | × | × | × | 7 | 85.42 | 6.46 | 0.018 |
| 19 | | × | | | × | 5 | 85.61 | 6.65 | 0.016 |
| 10 | × | | | × | | 5 | 85.89 | 6.93 | 0.014 |
| 17 | | | | | × | 4 | 85.93 | 6.97 | 0.014 |
| 18 | × | | | | × | 5 | 87.56 | 8.61 | 0.006 |
| 20 | × | × | | | × | 6 | 88.08 | 9.12 | 0.005 |
| 14 | × | | × | × | | 6 | 88.49 | 9.53 | 0.004 |

| | | | | | | | | | |
|---|---|---|---|---|---|---|---|---|---|
| 12 | × | × | | × | | 6 | 88.57 | 9.61 | 0.004 |
| 15 | | × | × | × | | 6 | 88.74 | 9.78 | 0.003 |
| 23 | | × | × | | × | 6 | 88.86 | 9.90 | 0.003 |
| 32 | × | × | × | × | × | 8 | 88.93 | 9.97 | 0.003 |
| 21 | | | × | | × | 5 | 88.96 | 10.00 | 0.003 |
| 3 | | × | | | | 4 | 89.39 | 10.43 | 0.002 |
| 1 | | | | | | 3 | 90.03 | 11.07 | 0.002 |
| 22 | × | | × | | × | 6 | 90.80 | 11.84 | 0.001 |
| 24 | × | × | × | | × | 7 | 91.57 | 12.61 | 0.001 |
| 2 | × | | | | | 4 | 91.67 | 12.71 | 0.001 |
| 4 | × | × | | | | 5 | 91.79 | 12.84 | 0.001 |
| 16 | × | × | × | × | | 7 | 92.18 | 13.22 | 0.001 |
| 7 | | × | × | | | 5 | 92.65 | 13.69 | 0.000 |
| 5 | | | × | | | 4 | 93.01 | 14.05 | 0.000 |
| 6 | × | | × | | | 5 | 94.76 | 15.80 | 0.000 |
| 8 | × | × | × | | | 6 | 95.27 | 16.31 | 0.000 |

**REFERENCES:**

Asghar, M., Westerdahl, H., Zehtindjiev, P., Ilieva, M., Hasselquist, D. and Bensch, S. 2012. Primary peak and chronic malaria infection levels are correlated in experimentally infected great reed warblers. - *Parasitology,* **139**: 1246-1252.

Bell, J. A., Weckstein, J. D., Fecchio, A. and Tkach, V. V. 2015a. A new real-time PCR protocol for detection of avian haemosporidians. - *Parasites & vectors,* **8**: 383.

Bell, J. A., Weckstein, J. D., Fecchio, A. and Tkach, V. V. 2015b. A new real-time PCR protocol for detection of avian haemosporidians. - *Parasites Vectors,* **8**: 9. https://doi.org/10.1186/s13071-015-0993-0

Bentz, S., Rigaud, T., Barroca, M., Martin-Laurent, F., Bru, D., Moreau, J. and Faivre, B. 2006. Sensitive measure of prevalence and parasitaemia of haemosporidia from European blackbird (*Turdus merula*) populations: value of PCR-RFLP and quantitative PCR. - *Parasitology,* **133**: 685-692.

Dawson, R. J., Gibbs, H. L., Hobson, K. A. and Yezerinac, S. M. 1997. Isolation of microsatellite DNA markers from a passerine bird, *Dendroica petechia* (the Yellow Warbler), and their use in population studies. - *Heredity,* **79**.

Formica, V. A. and Tuttle, E. M. 2009. Examining the social landscapes of alternative reproductive strategies. - *J. Evol. Biol.,* **22**: 2395-2408. https://doi.org/10.1111/j.1420-9101.2009.01855.x

Grunst, A. S., Grunst, M. L., Korody, M. L., Forrette, L. M., Gonser, R. A. and Tuttle, E. M. 2019. Extrapair mating and the strength of sexual selection: insights from a polymorphic species. - *Behavioral Ecology*.

Hall, T. A. 1999. BioEdit: a user-friendly biological sequence alignment editor and analysis program for Windows 95/98/NT. *Nucleic acids symposium series.* [London]: Information Retrieval Ltd., c1979-c2000., pp. 95-98.

Hellgren, O., Waldenström, J. and Bensch, S. 2004. A new PCR assay for simultaneous studies of *Leucocytozoon*, *Plasmodium*, and *Haemoproteus* from avian blood. - *Journal of Parasitology,* **90**: 797-802.

Jeffery, K., Keller, L., Arcese, P. and Bruford, M. 2001. The development of microsatellite loci in the song sparrow, *Melospiza melodia* (Aves) and genotyping errors associated with good quality DNA. - *Mol. Ecol. Notes,* **1**: 11-13.

Kalinowski, S. T., Taper, M. L. and Marshall, T. C. 2007. Revising how the computer program CERVUS accommodates genotyping error increases success in paternity assignment. - *Mol. Ecol.,* **16**: 1099-1106. https://doi.org/https://doi.org/10.1111/j.1365-294X.2007.03089.x

Peig, J. and Green, A. J. 2009. New perspectives for estimating body condition from mass/length data: the scaled mass index as an alternative method. - *Oikos,* **118**: 1883-1891. https://doi.org/https://doi.org/10.1111/j.1600-0706.2009.17643.x

Petren, K. 1998. Microsatellite primers from Geospiza fortis and cross-species amplification in Darwin's finches. - *Mol. Ecol.,* **7**: 1782-1784.

Pigeault, R., Cozzarolo, C.-S., Glaizot, O. and Christe, P. 2020. Effect of age, haemosporidian infection and body condition on pair composition and reproductive success in Great Tits *Parus major*. - *Ibis,* **162**: 613-626. https://doi.org/10.1111/ibi.12774

Poesel, A., Gibbs, H. L. and Nelson, D. A. 2009. Twenty-one novel microsatellite DNA loci isolated from the Puget Sound white-crowned sparrow, *Zonotrichia leucophrys pugetensis*. - *Mol. Ecol. Resour.,* **9**: 795-798.

Tuttle, E. M. 2003. Alternative reproductive strategies in the white-throated sparrow: behavioral and genetic evidence. - *Behav. Ecol.,* **14**: 425-432. https://doi.org/10.1093/beheco/14.3.425
